# Cross-excitation in dorsal root ganglia redistributes coding capacity from stimulus intensity to stimulus location

**DOI:** 10.64898/2026.11.05.764541

**Authors:** Dmitrii Perevozniuk, Pavel Musienko, Nikolay Koshev

**Affiliations:** Center for Bio- and Medical Technologies, Moscow, Russia; Laboratory of Neuroprosthetics, Institute of Translational Biomedicine, Saint Petersburg State University, Saint Petersburg, Russia; Neuroscience Department, Sirius University of Science and Technology, Sirius Federal Territory, Russia; Life Improvement by Future Technologies Institute (LIFT), Moscow, Russia; Moscow Center for Advanced Studies, 20, Kulakova Str., Moscow, Russia

**Keywords:** Dorsal root ganglia, Cross-excitation, Somatosensory processing, Neural coding, Mutual information, Leaky integrate-and-fire model

## Abstract

The phenomenon of cross-excitation between neurons of dorsal root ganglia (DRGs) of mammals is widely known, yet its exact mechanism remains underexplored. In this work, we analyze the phenomenon as part of the signal processing performed by the body’s somatosensory system. We argue that, if the mechanism is indeed as prevalent as it is reported in experimental works, it should provide a significant computational advantage. Here, we created a model network in which sensory A-type neurons are interconnected only via a chemical agent and showed that such a connection, when fitted with parameters similar to those observed in experiments, improves the stimulus-localization properties of the ganglia at the expense of intensity-encoding accuracy. Although this exchange is near-neutral in raw bits, it becomes favorable once the logarithmic compression of intensity perception — the Weber–Fechner law — is taken into account, since the sacrificed intensity capacity is largely perceptually redundant. We therefore propose that cross-excitation is not an imperfection of somatic isolation but an effective, chemically mediated pre-processing stage that sharpens spatial localization.

## 1 Introduction

Dorsal Root Ganglia (DRG) are specialized structures within the peripheral nervous system in which bodies of primary sensory neurons are located. From a histological perspective, these neurons are unique; they are pseudo-unipolar, possessing a single axon that emerges from the soma and bifurcates into two branches, forming a characteristic T-junction (Lawson 2005; Ross and Pawlina 2016). Furthermore, DRG somata are tightly enveloped by Satellite Glial Cells (SGCs), which form an envelope around the neuronal bodies (Pannese 1981; Hanani 2005; Matsuda et al. 2000). This glial ensheathment prevents the formation of synapses between neuron bodies. While the gaps in the envelope were reported, it remains unclear whether those are a property of this system or merely an experimental artifact (Shinder and Devor 1994; Pannese et al. 2003).

Physiologically, these neurons provide somatosensory functions — sending out endings of peripheral branch into skin and muscles, sensing a wide variety of modalities (such as pressure of different amplitudes, temperature sensing, etc.). Such neurons have been shown to have A-type fibers and sometimes are called large-light neurons (as they are 45 micrometers in diameter and appear light in microscope) (Shinder and Devor 1994; Lawson 2005). Another type of neurons — those having C-type fibers — are titled small dark neurons and carry nociceptive signals of various modalities (e.g., painful heat and cold, tissue damage, inflammation).

Each neuron corresponds to a specific region on the skin (or in a muscle), called a receptive field, where its dendrites end and to stimuli where it responds. Receptive fields vary in size, depending on the body part — the fields can range from several cm^2^ (as those located on the backside of human torso) and to less than mm^2^ (on fingertips). This property follows a clear evolutionary rationale: as fingertips are used to estimate various properties of object being touched (i.e. form, texture, temperature, etc.) they require much higher resolution, as opposed to sensors located on the back. Additionally, receptive fields are known to overlap (Lawson 2005), increasing resolution without increasing sensor density. Consequently, a higher number of receptive fields (and higher density of sensors) requires higher number of neuron bodies, which can be observed anatomically as an enlargement of cervical DRGs (brachial plexus), the ones housing neurons innervating palms. These properties represent a sensory adaptation to maximize information gain within limited resources. It is therefore logical to analyze this system as a technical solution to a signal detection problem.

From described anatomical isolation of neuron bodies (SGC envelopes) stems a view of the DRG as a passive relay station, where the primary function is simply to transmit signals from the periphery to the spinal cord, while filtering noise via thresholding. However, advanced experimental methods have challenged this view, revealing that some kind of a non-synaptic interaction exists. This phenomenon, known as cross-excitation (or cross-depolarization) (Devor and Wall 1990; Utzschneider et al. 1992; Shinder et al. 1998), occurs when depolarization of one neuron transiently lowers the excitation threshold (induces sub-threshold depolarization) in its neighbors.

The distribution of this phenomenon is intriguing. Cross-excitation is highly prevalent among large light (A-type) neurons, with over 90% exhibiting this interaction. In contrast, it is observed in less than 5% of small, dark (C-type) neurons (Devor and Wall 1990). This discrepancy probably arises from functional differences: while cross-talk may benefit mechanoreception or proprioception (mediated by A-fibers), it could be maladaptive in nociceptive C-fibers, where “smearing” pain signals across neurons could lead to pathological conditions such as hyperalgesia (Dublin and Hanani 2007; Spray and Hanani 2019; Zhang et al. 2009).

While the precise biophysical mechanisms — ranging from neurotransmitter leakage through SGC gaps (Rozanski et al. 2012, 2013a; Kung et al. 2013; Mandge et al. 2017; Rozanski et al. 2013b) to transient potassium accumulation (Mandge et al. 2019; Sibille et al. 2015; Suadicani et al. 2010) — remain a subject of debate, the existence of cross-excitation is well-established (Devor and Wall 1990; Shinder et al. 1998; Utzschneider et al. 1992; Huang et al. 2006). This leads to a fundamental question regarding the evolutionary utility of the DRG cross-excitation. If this interaction is so prevalent among specific sensory types, does it provide a functional advantage?

To properly discuss the DRG as a signal detecting and preprocessing tool, the principles of neural coding should be addressed. Sensitive endings (sensors of various modalities) produce two types of potentials — action potentials (which are uniform in amplitude and shape) and transient generator potentials (essentially — subthreshold shifts of membrane potential). While the former ones can traverse the whole length of nerve after being generated at an ending, the latter can not travel far on their own. When enough transient potentials are summed (at the spike initiation zone) a thresh-old is overcome and an action potential is formed. However, transient potentials can arise due to some other influences (either random molecular fluctuations of sensor states or an effect of forces of other modalities) so this analogous encoding can also be considered a source of noise. On the other hand, the action potentials (generated by sensitive endings when the excitation threshold is surpassed) encode the information about the presence and intensity of a particular stimulus. On the periphery, rate coding is the primary mode of transmission — while the stimulus causes the endings to spike, the intensity of it (i.e., applied pressure or temperature) affects the spiking frequency. Further, sensory neurons can be divided into two categories — fast and slow adapting, based on their adaptation rates. Fast adapting (FA) neurons fire only at the onset and offset of a stimulus, effectively encoding the rate of change (velocity) rather than absolute intensity. Conversely, slowly adapting (SA) neurons maintain firing throughout the stimulus duration, proportionally encoding the static amplitude. Finally, the spatial properties (where the stimulus is applied) are encoded implicitly — the neurons responsible for nearby receptive fields are neighboring in DRG, as well as their filaments arriving into one structure side-by-side. This idea, a clear correspondence between structural (where the neurons and their endings are located) and functional (what areas do they sense) is called a somatotopy, and it can be observed in various sensory systems (proprioception, vibrissae sensing, visual — there it would be called retinotopy, etc.) of all animals (Kandel et al. 1991). Thus, the afferent signal arriving from the DRG is a multidimensional vector containing the spatial position, intensity, and the first time derivative of intensity of the external stimulus.

In the present work we investigate the physiological implications of cross-excitation existing. We ask: What does the nervous system gain from the coupling of somatosensory neurons? Is cross-excitation merely an “imperfection of isolation” — a biological artifact of tightly packed somata — or does it represent a sophisticated mechanism for sensory information preprocessing? By simulating the DRG network under varying conditions, we aim to determine if this phenomenon enhances signal fidelity, synchronization, or noise robustness, thereby proving the DRG as an active information processing center rather than a passive pre-amplifier.

## 2 Materials and Methods

### 2.1 Modeling and simulations

All described simulations were scripted using Python programming language, with wide usage of such libraries as NumPy (Harris et al. 2020), SciPy (Virtanen et al. 2020), Scikit (Pedregosa et al. 2011), matplotlib (Hunter 2007), Numba (Lam et al. 2015).

#### 2.1.1 Overall architecture

To test the hypothesis of cross-excitation improving somatosensory system’s properties in terms of spatiotemporal stimulus detection, we needed a model of the system and a way to measure its performance.

We recreated a leaky integrate-and-fire (LIF) neural network (Abbott 1999), in which neurons lay side by side without any connections to peers, with resting potentials initiated randomly for each neuron in range from *−*75 mV to *−*55 mV (Davidson et al. 2014). No synaptic connections are established, as this structure normally lacks any synapses between its neurons (Lawson 2005). Activation thresholds were also randomized between +12.5 mV and +37.5 mV from resting potential (Ho and O’Leary 2011). Both distributions are uniform within the stated boundaries. With this, we aimed to achieve more biologically plausible activity, as neurons with such an approach tend to fire in characteristic batches and not simultaneously on the same simulation step (Fig 1). This is due to the fact that, in a real system, the excitability threshold of a particular neuron type is not, in fact, a single value, but rather a distribution in some range.

**Fig. 1.**
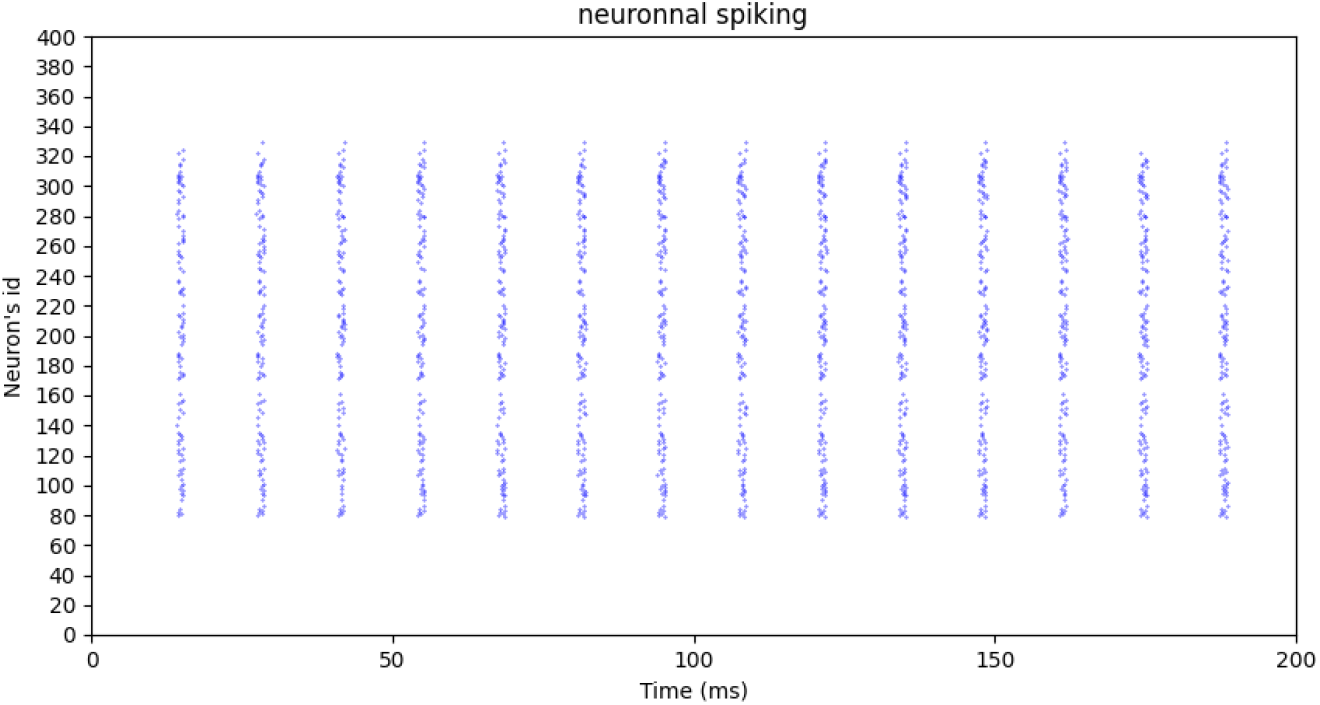
Example of neuronal spiking in our model. Note the characteristic batches of spikes, which arise due to differences in excitation thresholds.

Absolute refractory periods of 2 ms were introduced for each neuron. LIF model was chosen as a balance between biological plausibility and computational efficiency. It allows one to simulate networks composed of thousands of spiking neurons by abstaining from directly calculating ionic fluxes and geometrical properties of the cells, as opposed to more classical Hodgkin-Huxley formalism (Hodgkin and Huxley 1952).

As for the estimation of the network performance, we introduced a slightly modified intersection over union (IOU) metric (IOU) (Everingham et al. 2010). Our time-spatial metric evaluates the accuracy of neural responses through two distinct Jaccard index components: temporal and spatial. The temporal component assesses event-level detection by separating the timeline into stimulation and gap periods based on the actual presence of an input stimulus. A true positive is recorded if the neuron spikes during a stimulation period (and if stimulus is located in its receptive field, i.e., it indeed receives the stimulating current), a false negative if it remains silent during stimulation, and a false positive if it spikes during a gap. The spatial component evaluates the precision of the population response within each stimulation period. For every neuron, a ground truth activation is defined by its cumulative input exceeding the power threshold, while its predicted activation is defined by the presence of at least one spike during that stimulation window. Both components are calculated using the intersection over union formula, dividing the total number of true positives by the sum of true positives, false positives, and false negatives. Two components are either analyzed separately or the mean is calculated, to produce a single spatio-temporal IOU metric, which characterizes spatio-temporal accuracy of the system.

Neurons had only one input and one output and were not connected to each other by synapses, thus imitating isolated neurons of DRG. To approach biological plausibility, we imitated transient generator potentials by randomly applying subthreshold noise to neurons’ inputs.

Additionally, we implemented a fiber recruitment ratio (Dalrymple et al. 2021) — a neuron which is actually stimulated can, with a particular probability, be mistaken and not emit a spike.

Further, the current applied was always equal to current delivered by 75 mV stimulation over 10 ms, no matter the stimulus length. This amplitude was picked as it is sub-threshold for half of the neurons generated as described above.

We introduced a chemical agent into the system — since the actual parameters of such an agent (if we follow the chemical mediator hypothesis of cross-excitation) is not known, we needed to fit this from reported experimental observations. We modeled the dynamics of said agent as a simple exponential decaying concentration (Syková and Nicholson 2008), where the agent effect on neurons is calculated as the weight *g*_*ce*_ multiplied by concentration of it near the neuron. When the neuron fires, it adds a particular amount of agent to all connected neurons.

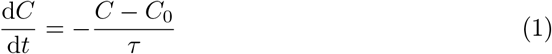

Here *C* is the current concentration of the agent (mM), *C*_0_ the initial concentration (set to 0 mM), *τ* the degradation constant (ms) and *t* the time (ms).

The degradation constant *τ* remains unknown, as we do not know actual properties of the molecule. However, we can simply fit it, together with *g*_*ce*_ coefficient: the CE is known (Devor and Wall 1990; Utzschneider et al. 1992) to shift membrane potential of the affected neuron within half a second for *≈*+10 mV. Simulating one neuron whose neighbor is affected by 50 Hz stimulation and analyzing at which values of *τ* and *g*_*ce*_ the membrane potential crosses said threshold in 500 ms (thus imitating the original experiment from (Devor and Wall 1990)). We searched for parameters that suffice and obtained approximate values for said constants, *τ* = 220 ms and *g*_*ce*_ = 0.22.

Thus, the total equation governing the membrane potential of each neuron was

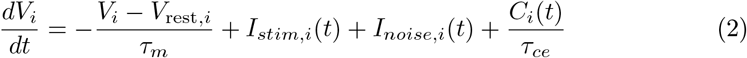

All simulations were performed on a population of *N* = 400 neurons, for *T* = 2000 ms of simulation time, with *dt* = 0.05 ms.

Since the equations above assume an immediate and equal deposit to all of the neighbors, this model can also be interpreted as a model of interaction between immediate neighbors – in this case, the delivery time would be minuscule and the proportions approximately equal. As the phenomenon of cross-excitation shall yield different results under different conditions, we ran several simulations, analyzing timespatial accuracy of the model in various setups. Additionally, we decided to compare two situations: the first — ideal scenario — is a model with low noise and high afferent recruitment. The second — a “noisy” scenario — featuring high noise and significant stochastic input failure. This variety of setups also helps us to generalize CE effects despite high biological variability. Even if we do not know the exact properties of said interaction (and even some properties of the system itself — i.e., noise probability and distribution), we hope to derive general conclusions that will hold true in most cases. Since the model contains a stochastic part, each point was estimated 5 times, and the results are given as mean *±*SD.

Note that in graphs CE range is reported as neurons to 1 side. It is a simplification, which is tied to the way the model was programmed and it assumes 1D structure, in which a neuron interacts with *N* neighbors on both sides (which, assuming symmetry of the effect, is reported as a CE range of *N* neurons to one side).

For each experiment, we also calculated relative mutual information heatmaps. These were calculated as the mutual information between the output and input of the system with cross-excitation minus the mutual information of the same system under the same stimulation conditions without cross-excitation. This way, we observed that although simple IOU metrics show an insignificant increase, the overall net gain in information is still (slightly) positive in most cases (see the corresponding graphs).

### 2.2 Theoretical derivations

The simulations show a consistent trade: cross-excitation lowers temporal accuracy yet raises spatial accuracy, with a small net gain in mutual information (see the simulation results below). Here we present theoretical explanations for this trade from the model equations. The argument proceeds in four steps. We first describe the CE-free ganglion as a group of independent noisy channels (*Baseline*). We then derive the dynamics of the diffusible agent that couples neurons, showing that its mean drive grows linearly with firing rate and switches on above a definite frequency (*Cross-excitation coupling* ). Next we show that the resulting spatial gain scales with the unreliability of the baseline code (*Spatial enhancement* ). Finally we assemble these results into a single information-theoretic statement (*Information-capacity redistribution*): CE does actively reallocate the population’s finite coding capacity from stimulus intensity to stimulus location, at a fixed cost of a few bits, and this reallocation is favorable once the perceptual value of the two channels – set by the Weber–Fechner law – is taken into account.

#### 2.2.1 Baseline: the ganglion as a set of parallel noisy channels

Without CE, the neurons of Dorsal Root Ganglion are uncoupled: each obeys its own leaky integrate-and-fire (LIF) equation and receives only its afferent drive and independent noise. A single neuron is therefore a noisy binary channel – it either does or does not emit a spike in a given read-out window – and, because the channels are independent, the information the population carries is the sum of the per-neuron contributions, i.e., the ganglion can be represented as the sum of independent noisy channels (Borst and Theunissen 1999; Stone 2018). This is the reference against which every CE effect in this paper (relative IOU, relative mutual information) is measured. Under rate coding (Johnson 2001) the amplitude *A* is written into the firing rate, *f ∝A*, and read out as a spike count over a window *T*. Modeling the count as an approximately Gaussian variable whose mean and variance both scale with the number of spikes, the intensity channel has a signal-to-noise ratio set by the expected count,

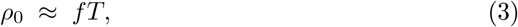

so the baseline information the response carries about intensity, at a fixed location, is the Gaussian-channel value

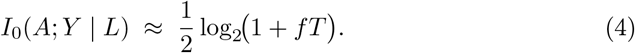

Eqs (3) and (4) say that, without CE, more stimulation (higher *f* ) buys more intensity information without limit. We return to this unbounded linear growth of raw capacity below: it is the surplus that CE is able to spend.

#### 2.2.2 Cross-excitation coupling and agent dynamics

We model CE as a diffusible agent whose local concentration *C*_*i*_(*t*) adds a depolarizing term *C*_*i*_(*t*)*/τ*_*ce*_ to the membrane equation of neuron *i*, Eq (2). Each neighbor spike deposits a fixed quantum *g*_*ce*_ of agent that then decays exponentially with time constant *τ*.

##### Steady-state accumulation

Consider one neighbour firing at a constant rate *f*, i.e. one spike every Δ*t* = 1*/f*. The *k*-th preceding spike was emitted a time *k/f* ago and its residual has decayed to *g*_*ce*_ *e*^*−k/*(*fτ*)^. Summing the geometric series of all past spikes gives the inter-spike peak concentration,

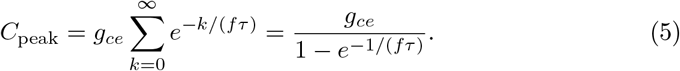

Averaging the decaying concentration over one inter-spike interval collapses to a strikingly simple result,

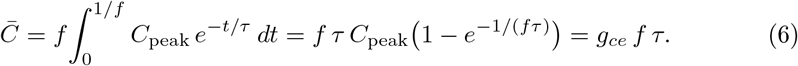

For *N*_*k*_ active somatotopic neighbors firing at the common rate *f* the mean drive simply adds,

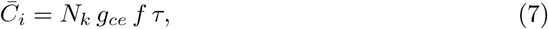

so the average agent concentration — and hence the mean CE depolarization — is *linear in both the firing rate and the number of coupled neighbours*. This linearity is the hinge of every argument that follows. **The functional threshold** *f*_th_. Accumulation only becomes *functional* once the mean CE drive can bridge the gap between rest and threshold. For a constant drive the steady-state offset of the LIF equation (2) is 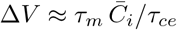, so CE lifts the membrane by

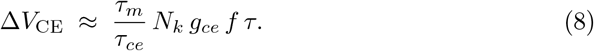

Setting this equal to the mean gap between resting and excitation potential, Δ*V*_gap_ *≈*25 mV, defines the functional threshold

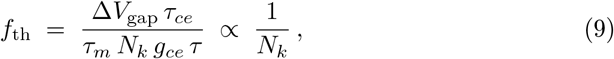

which for the fitted parameters and *N*_*k*_ = 7 evaluates to *f*_th_ *≈*24.3 Hz. Two features of (9) recur later: *f*_th_ is a genuine switch-on frequency (below it CE does little), and it falls as the interaction range *N*_*k*_ grows.

##### Three regimes

Equation (5) distinguishes three regimes of periodic drive, separated by the product *fτ*:

- **Sub-accumulation** (*fτ ≪* 1): spikes arrive far apart, each residual decays before the next, *C*_peak_ *→g*_*ce*_ and the time-averaged drive (6) is negligible.
- **Accumulation onset** (*fτ* ≳ 1): residuals of consecutive spikes begin to overlap. With the fitted *τ ≈* 220 ms this transition sits at *f* = 1*/τ ≈*4.5 Hz; the cumulative tail is still small and saturates quickly because decay is exponential.
- **Functionally effective** (*f* ≳ *f*_th_): the accumulated drive from all *N*_*k*_ neighbors is large enough to matter dynamically (below).

#### 2.2.3 Information-capacity redistribution

We now assemble the preceding results into a single information-theoretic statement. Driven by a stimulus *X* = (*L, A*) and returning the binary spike vector *Y* = (*Y*_1_, …, *Y*_*N*_ ), the population’s response carries information that, by the chain rule of mutual information (Cover and Thomas 2005), splits into a spatial and an intensity term,

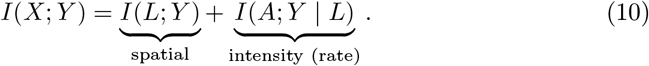

The first term measures how well *Y* identifies where the stimulus is; the second – how strong it is, given the location. These map directly onto our two empirical metrics: the spatial-domain IOU tracks *I*(*L*; *Y* ) and the temporal-domain IOU tracks *I*(*A*; *Y* | *L*). Our claim is that CE increases *I*(*L*; *Y* ) at the expense of *I*(*A*; *Y* | *L*), and that the trade is worthwhile.

#### A frequency-independent temporal cost

Without CE the intensity channel has the SNR *ρ*_0_ *≈ fT* of Eq (3). Switching CE on replaces the clean drive with the coupling term *C*_*i*_(*t*)*/τ*_*ce*_, whose value depends on the noisy firing of the neighbors; even with the stimulus *X* known exactly, *Y*_*i*_ therefore inherits residual uncertainty. Because *C*_*i*_(*t*) linearly depends on the neighbors’ spike trains, its variance is linear in *f*. Counting spike opportunities as independent Bernoulli events (Brillinger 1988) with per-window variance Var(*Y*_*j*_ | *X*_*j*_) = *p*(1 −*p*) and summing over the *N*_*k*_ neighbours firing at the common rate *f*,

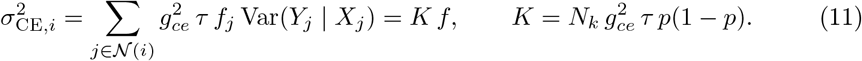

Crucially 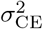 grows in *f in exactly the same proportion* as the mean drive 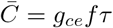 of Eq (6) The effective rate SNR under CE is therefore

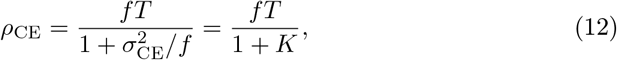

and the frequency dependence cancels. Using the Gaussian-channel form 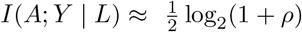, (Shannon 1948; Cover and Thomas 2005) the CE-induced change is

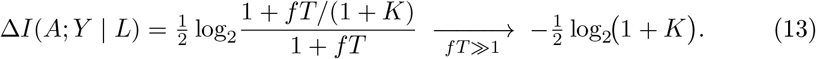

The temporal cost of CE is thus, to leading order, a *fixed number of bits*, independent of stimulation frequency: raising *f* amplifies the rate signal and the CE-produced noise in equal measure. Its magnitude is set entirely by the coupling constant *K* – i.e. by the CE calibration (*g*_*ce*_, *τ, N*_*k*_) – and not by the stimulus. This is exactly the temporaldomain pattern we observe (Fig 6, Fig 7): CE depresses temporal accuracy uniformly across the frequency axis rather than in any particular band, and the depression deepens with the interaction range *N*_*k*_, which enters *K* linearly (11), rather than with *f*.

#### A regime-dependent spatial gain

The spatial term behaves entirely differently. From the gap-filling result, the recovered firing that CE injects into the population image, weighted by how well that image is spatially organized, yields a spatial-information gain

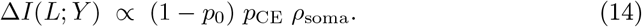

Equation (14) must *not* be read as tracking the mean drive 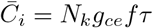 and therefore growing without bound in *f*. Its two rate-dependent factors pull in opposite directions. The gap-filling probability *p*_CE_(*f* ) rises with 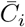, switching on near *f*_th_ (9) and saturating toward 1 thereafter. The available room for improvement (1 *−p*_0_(*f* )), by contrast, shrinks with *f*: more stimulation events per window make the baseline code more reliable, leaving fewer gaps to fill. Their product is therefore single-peaked — CE helps most in the band where the baseline is still leaky enough to leave gaps yet the neighbor drive is already dense enough to bridge them — and decays at high *f* where the code is near-saturated. This matches Fig 8 and the noisy spatial map Fig 9, in which spatial gains concentrate at lower frequencies and fade toward the high-frequency edge.

The same factor explains the noise dependence: under noise *p*_0_ is small, (1 *−p*_0_) is large, and the maximal spatial gain ( *≈* 0.226) far exceeds the ideal-case value ( *≈* 0.12), where the code is already reliable and there is little to recover. Finally, disrupting somatotopy lowers *ρ*_soma_ and, through (14), monotonically erodes the spatial gain — exactly the trend of the degree-of-distortion sweep (Fig 24).

#### The net exchange

Combining the two terms, the net information change from CE is

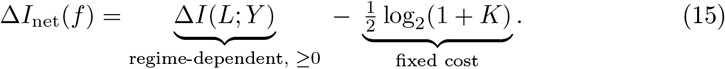

The raw balance is positive wherever gap-filling dominates: low-to-moderate frequency, modest interaction range, and especially the noisy regime; and turns mildly negative where the fixed intensity cost dominates, at high frequency and large range where the baseline is already reliable. This is precisely the structure of the relative mutual-information map (Fig 10): a positive effect at low-to-moderate frequency and small CE range, shading to slightly negative values at high frequency and large range. On a bit-for-bit accounting, then, CE is close to neutral, with a small positive balance in most conditions — consistent with the modest net-MI increase we report.

#### Why the exchange is favorable: excess bandwidth and the Weber–Fechner law

A bit-for-bit accounting treats a spatial bit and an intensity bit as equally valuable. They are not, and the intensity bits CE sacrifices turn out to be, perceptually, largely excessive.

In a rate code the firing rate *f* plays two roles at once: it is the signal (*f∝ A*) and it is the bandwidth, since a neuron emitting *f* spikes in a window *T* offers up to

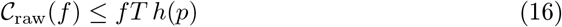

binary coding events (with *h*(*p*) the per-spike binary entropy), growing linearly with *f*: the same unbounded growth already visible in the baseline capacity (4). How much of this does intensity coding actually require? By the Weber–Fechner law perceived intensity is compressive (Portugal and Svaiter 2010),

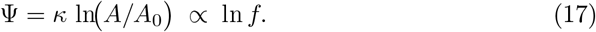

Equivalently, the just-noticeable difference is a fixed fraction of amplitude (Δ*A/A* = const), so the number of perceptually distinguishable intensity levels (*M* (*f* )) between the reference *A*_0_ and *A* is growing only logarithmically. The fraction of the linearly growing raw capacity that intensity discrimination needs is therefore

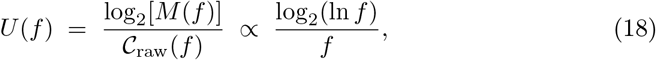

which falls monotonically for *f > e*. Thus the capacity that can be effectively utilized for intensity discrimination at 200 Hz is only *U* (200 Hz)*/U* (10 Hz) *≈*10% of that at 10 Hz. As the stimulus strengthens, an ever-larger share of the rate channel becomes redundant for perceptually useful intensity coding, and the surplus *C*_raw_ *−M ∝ f −*In *f* grows essentially linearly.

Two consequences follow. First, the fixed temporal cost of CE (13) is drawn from this increasingly redundant capacity: the same ½ log_2_(1+*K*) bits that are perceptually significant near threshold become an ever-smaller fraction of an ever-larger surplus at high *f*, so their marginal perceptual value *declines* with stimulus strength. Second, spatial localization is perceived as it is, without logarithmic compression, so converting surplus rate capacity into spatial information yields perceptual returns that do not diminish in the same way. The exchange that looks close to break-even in raw bits (15) is therefore favorable once weighted by perceptual value: CE spends intensity capacity that is cheap and getting cheaper to buy spatial capacity that stays valuable.

This resolves the apparent tension between the two frequency trends. The perceived payoff of CE is largest at low-to-moderate frequency and under noise, where real gaps exist to be filled (Fig 9); at high frequency the direct spatial payoff is small, but so is the perceptual cost of the intensity it mildly corrupts (18), so CE remains at worst harmless. Across the whole operating range the mechanism either helps materially or does no meaningful perceptual damage. Therefore the exchange is perceptually near-costless and brings extra bits of gain to spatial localization, i.e., favorable.

#### Summary of theoretical part

Cross-excitation acts as an active, non-synaptic spatial pre-filter that redistributes coding capacity along three mutually reinforcing lines. Its temporal cost is a fixed, frequency-independent number of bits (13), set by the coupling constant *K* rather than by the stimulus. Its spatial benefit scales with channel unreliability through the factor (1 *−p*_0_) (14), making CE most effective exactly when the system is stressed. And its benefit is gated by somatotopic coherence *ρ*_soma_, so the coupling injects structured rather than random perturbation. Because rate coding leaves a linearly growing surplus of capacity that perception, being logarithmic, cannot use (18), the net reallocation from intensity to location is perceptually adaptive rather than merely information-neutral — CE is a clever, chemically mediated preprocessing stage, not an imperfection of isolation.

## 3 Simulation results

### 3.1 Experiment 1

First, we tested the system under a wide variety of stimulation frequencies (2-200 Hz) and with varying CE range. Additionally, two scenarios were tested. The ideal scenario included noise probability (a chance for each neuron on each step to be affected by noise of amplitude 14% of stimulus) of 0.05 and fiber recruitment ratio of 1. The noisy scenario included noise probability of 0.25 and recruitment ratio of 0.75. Interestingly, presence of cross-excitation seems to decrease accuracy in time-domain (temporal accuracy of response) while increasing spatial accuracy. Additionally, the mean gain through all frequencies and CE ranges was higher in the noisy scenario (Fig 2, Fig 3, Fig 4, Fig 5). From this, we can also derive optimal cross-excitation range which increases time-spatial accuracy for most stimulation frequencies. Interestingly, the gain in spatio-temporal accuracy in both ideal and noisy scenario is insignificant (maximum gain in accuracy 0.022 for the ideal and 0.00115 for the noisy scenario). This, accompanied by optimal range of CE being 1 affected neighbor to both sides for noisy and 3 for ideal conditions proves that presence of cross-excitation does not, in fact, improve overall accuracy significantly. However, analyzing time and spatial accuracy separately, we observe an intriguing discrepancy: while the effect decreases or does not affect the temporal accuracy (maximum accuracy gain *≈* 0 ( −2.59*×*10^*−*6^ 1.13*×*10^*−*5^ for ideal and 0.0002 *±*0.00006 for noisy conditions; Fig 6, Fig 7), it greatly enhances the spatial accuracy (maximum gains of *≈*0.12 *±*0.091 and *≈*0.2264*±*0.051; Fig 8, Fig 9). Worth noticing that the increase in spatial gain is also much more pronounced in the noisy scenario. Optimal CE range, if optimized for maximum spatial accuracy, was 13 for the poor conditions and 7 for the ideal conditions.

**Fig. 2.**
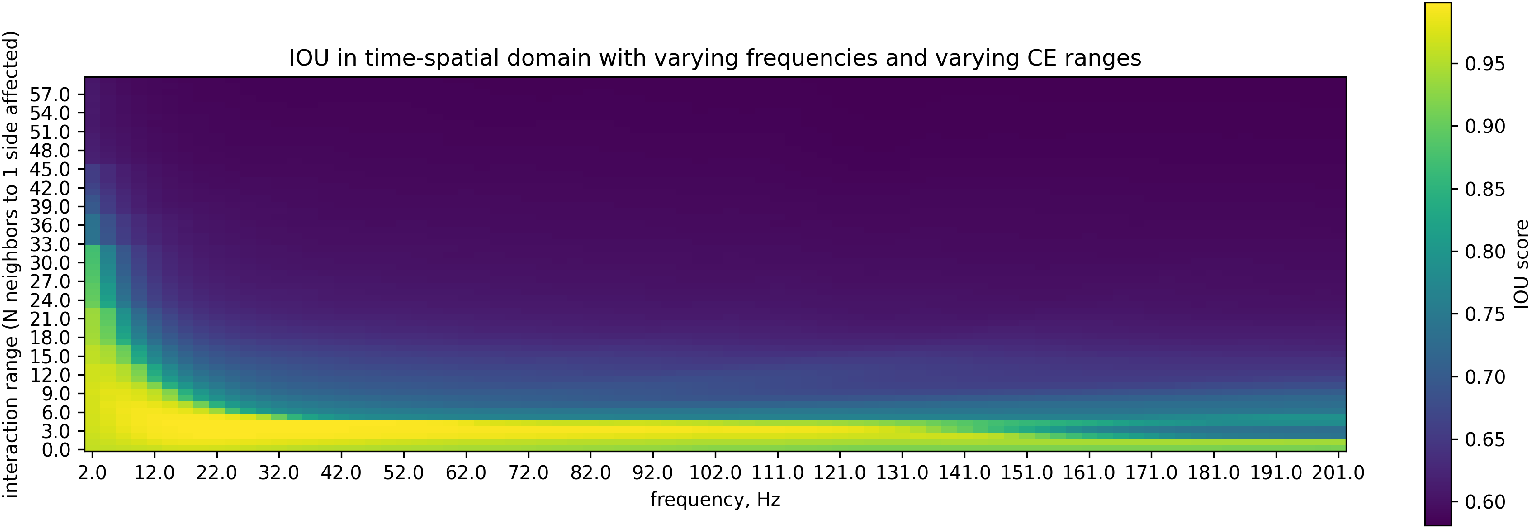
Accuracy of the system in time-spatial domain under ideal conditions.

**Fig. 3.**
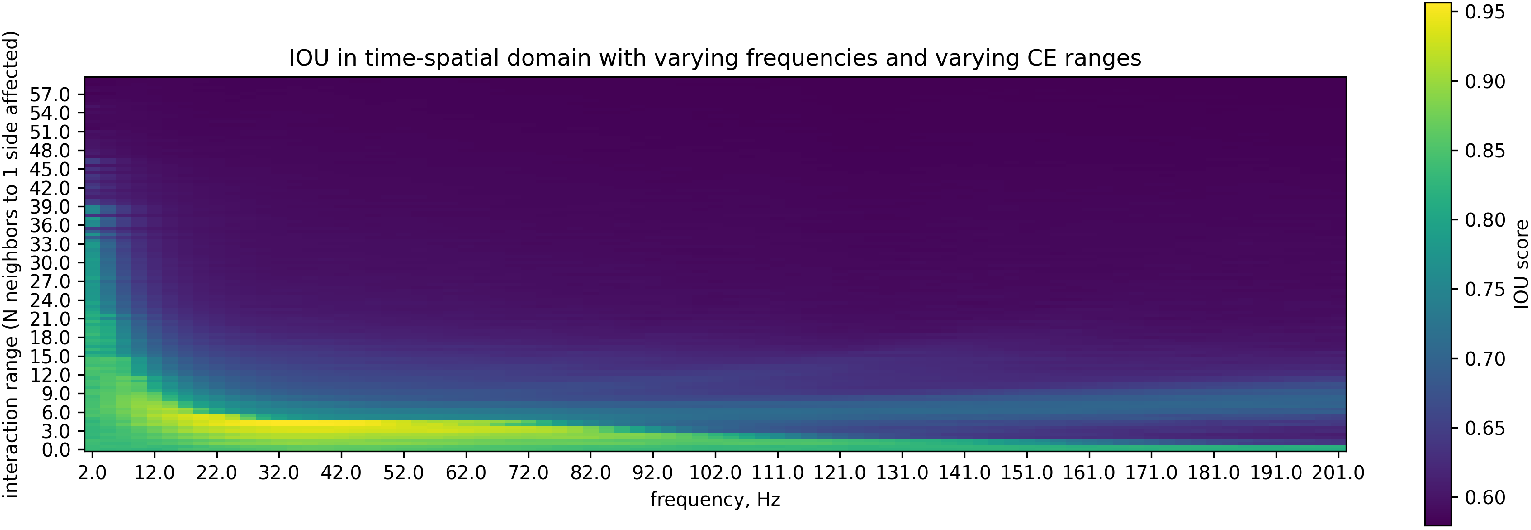
Accuracy of the system in time-spatial domain under noisy conditions.

**Fig. 4.**
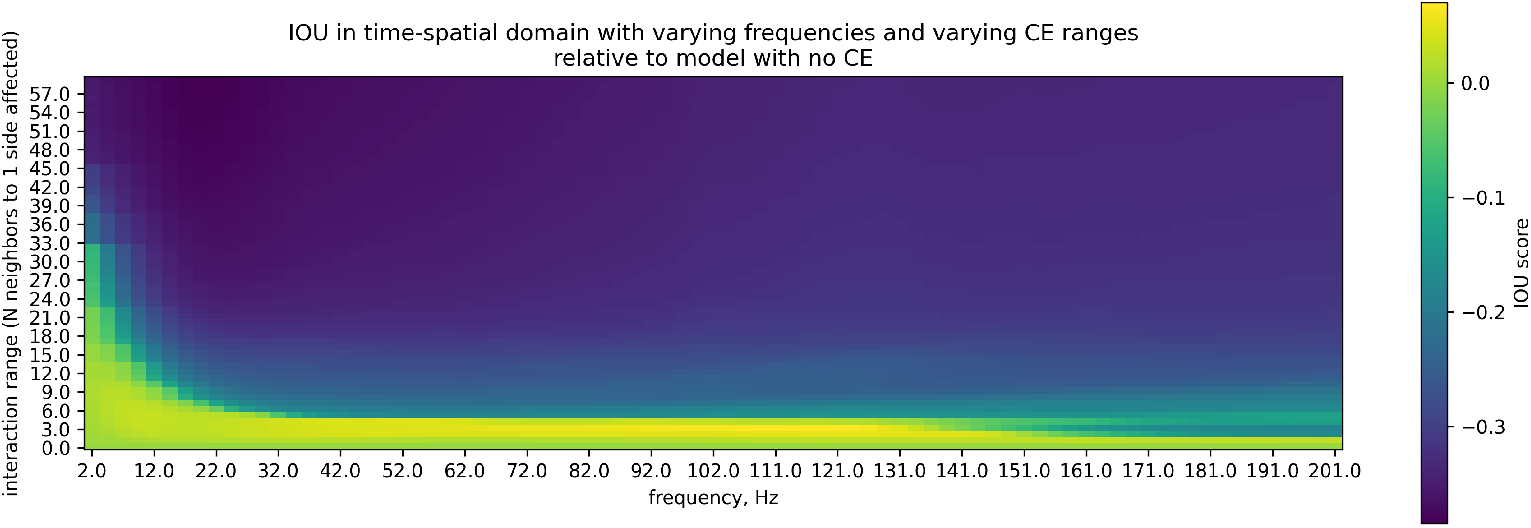
Relative accuracy of the system in time-spatial domain under ideal conditions.

**Fig. 5.**
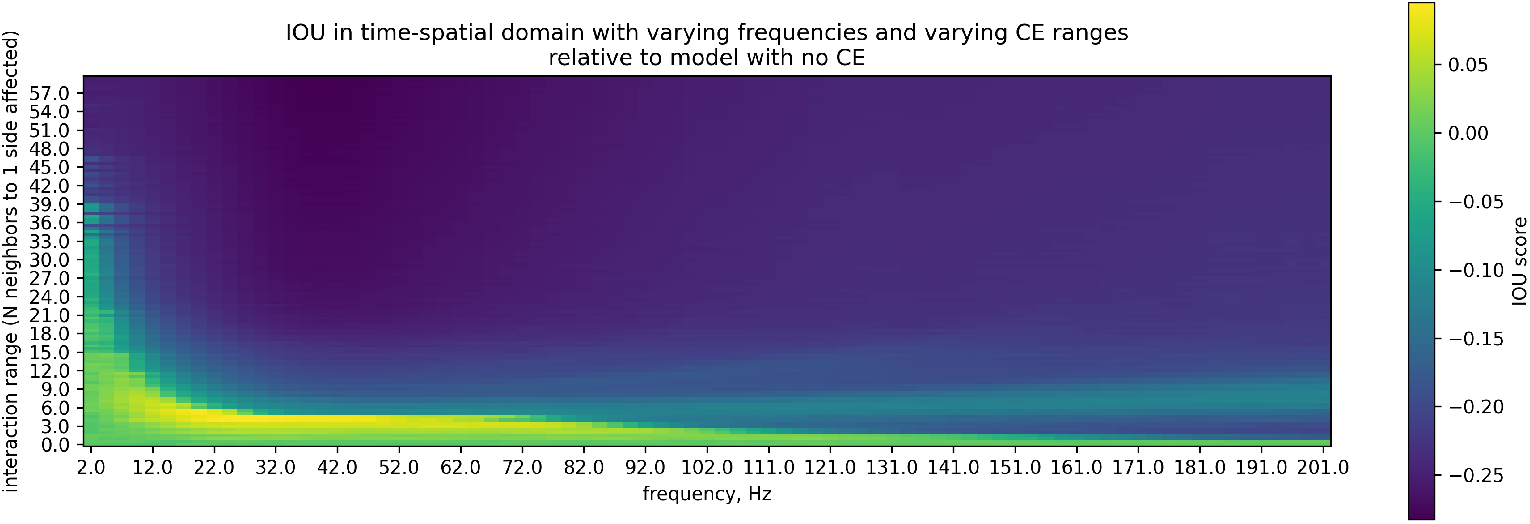
Relative accuracy of the system in time-spatial domain under noisy conditions.

**Fig. 6.**
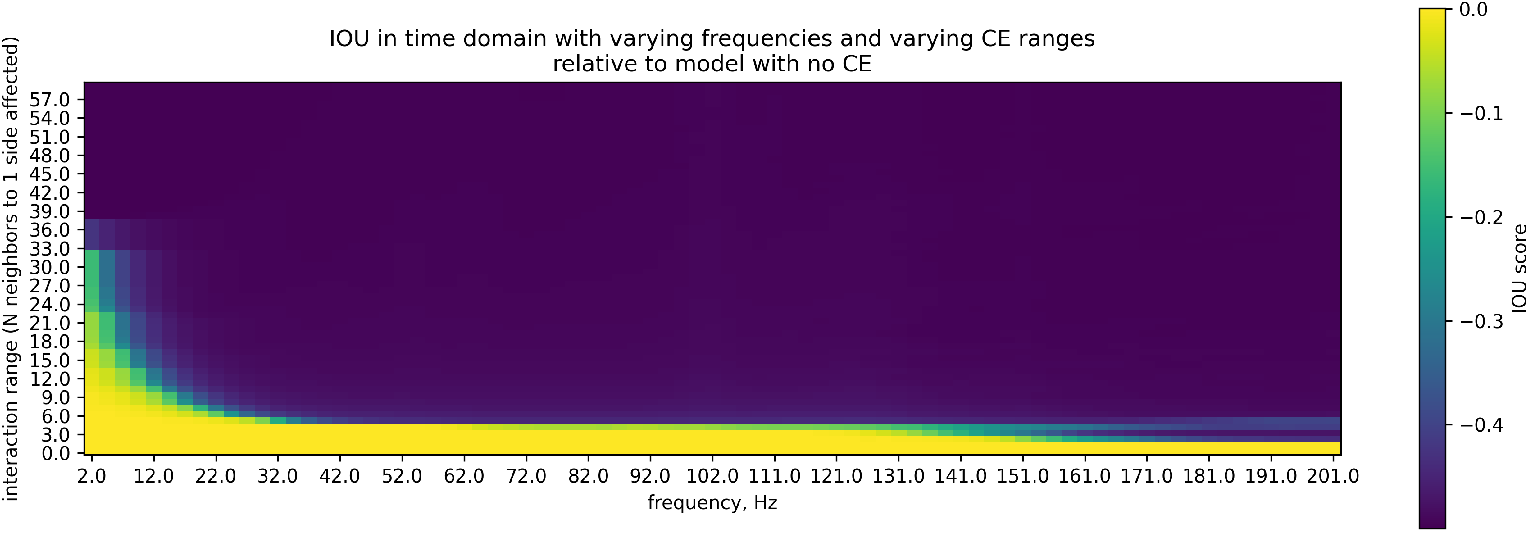
Relative accuracy of the system in time domain under ideal conditions.

**Fig. 7.**
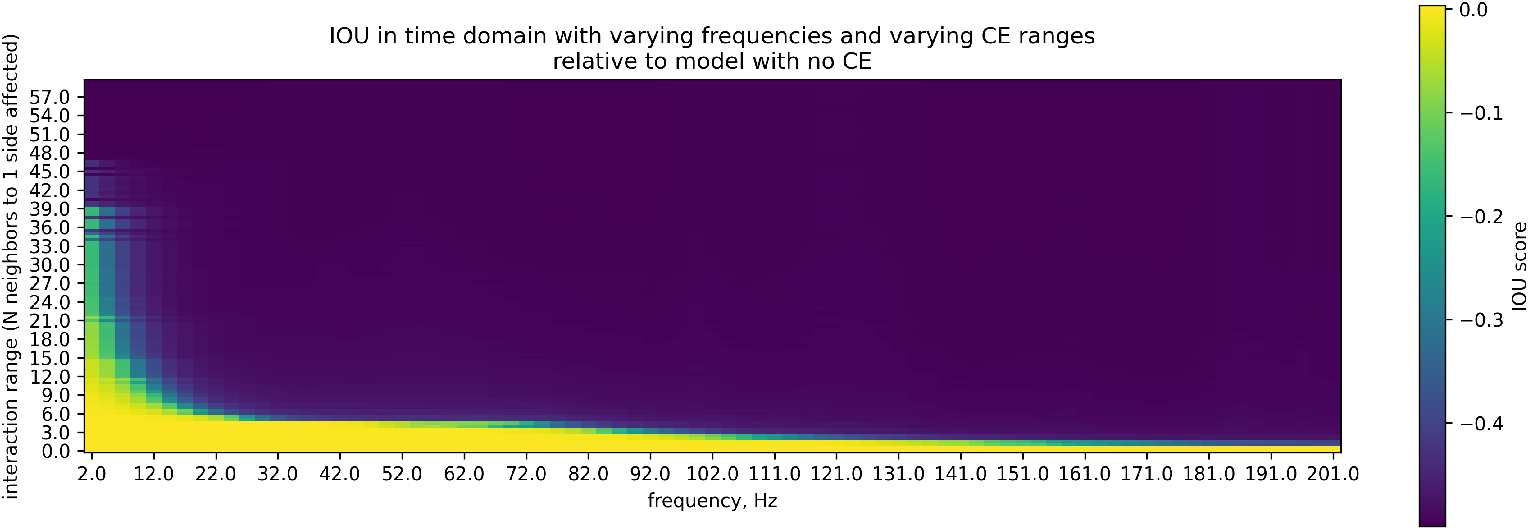
Relative accuracy of the system in time domain under noisy conditions.

**Fig. 8.**
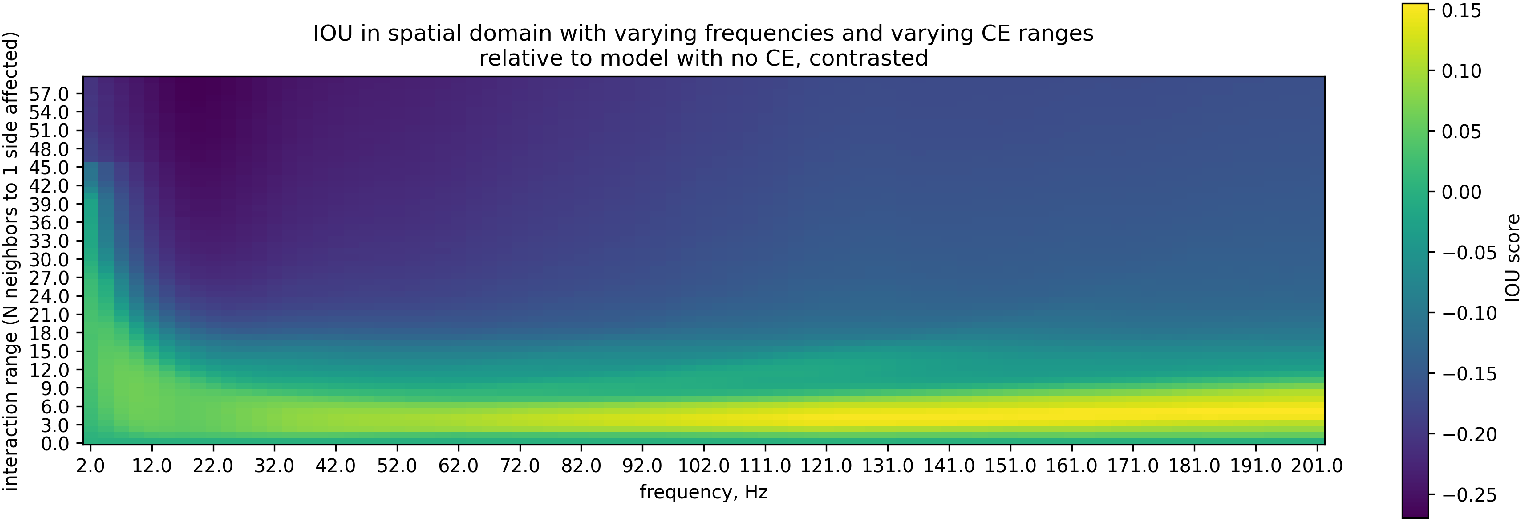
Relative accuracy of the system in spatial domain under ideal conditions.

**Fig. 9.**
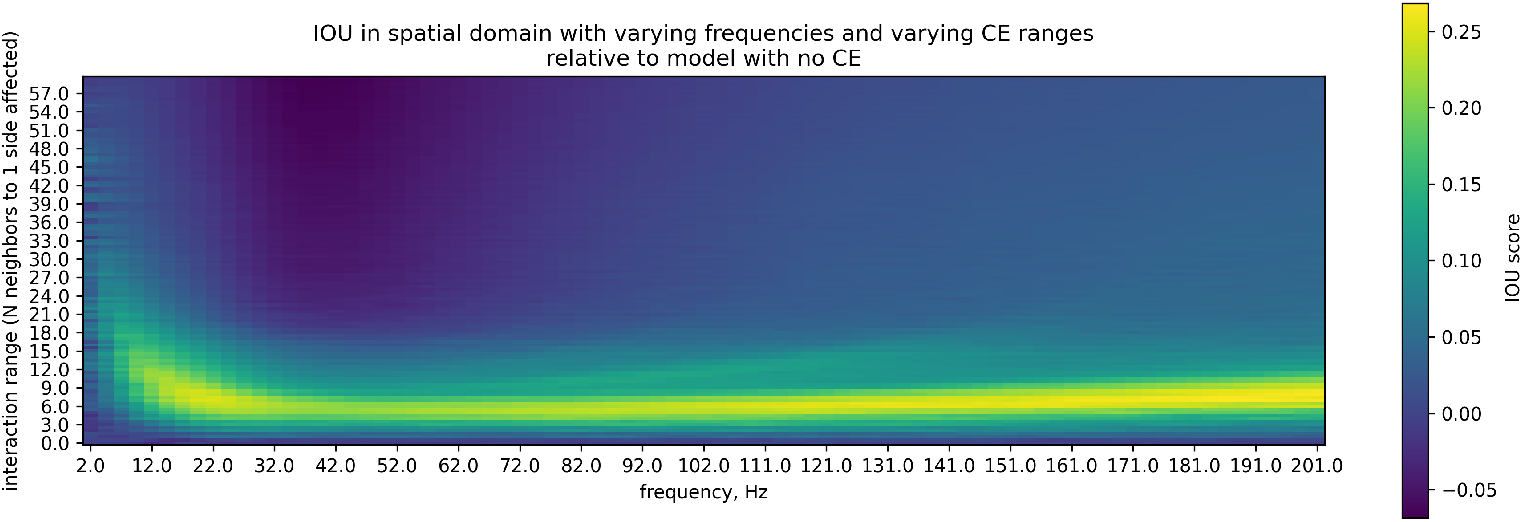
Relative accuracy of the system in spatial domain under noisy conditions.

**Fig. 10.**
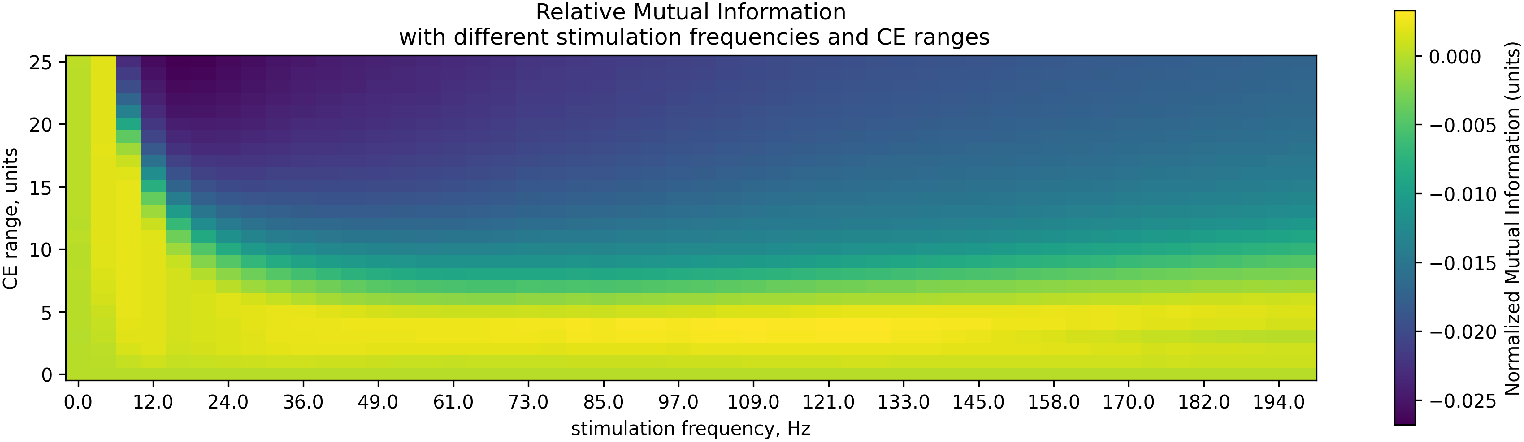
Relative mutual information under different stimulation conditions and varying CE range.

### 3.2 Experiment 2

Second, we tested a system with different noise parameters, as actual noise distributions of somatosensory nerves are not known. The noise was generated as a random input of fixed amplitude applied to a neuron on each simulation step with fixed probability. Here, the model was run in the noisy scenario and with a CE range of 7. We varied both amplitude and occurrence (probability) of the noise. Hence, the distribution of the noise in time and space was uniform, while its power (signal to noise ratio) varied. In this simulation, we systematically varied noise probability and frequency and calculated the relative IOU score (against the system without cross-excitation; Fig 11). Similarly, noise amplitude was varied together with frequency (Fig 14). The overall gain from CE was positive in the spatial domain (median gain through all frequencies and noise amplitudes *≈* 0.0454 *±* 0.0071; Fig 16) and mostly negative in the time domain (gain *≈ −*0.0555 *±* 0.0089; Fig 15). Similarly, through all combinations of noise occurrence probabilities and frequencies, the gain in the time domain was negative (mean *≈ −*0.12 *±* 0.084; Fig 12), while the gain in the spatial domain was positive (median *≈* 0.054 *±* 0.0093; Fig 13).

**Fig. 11.**
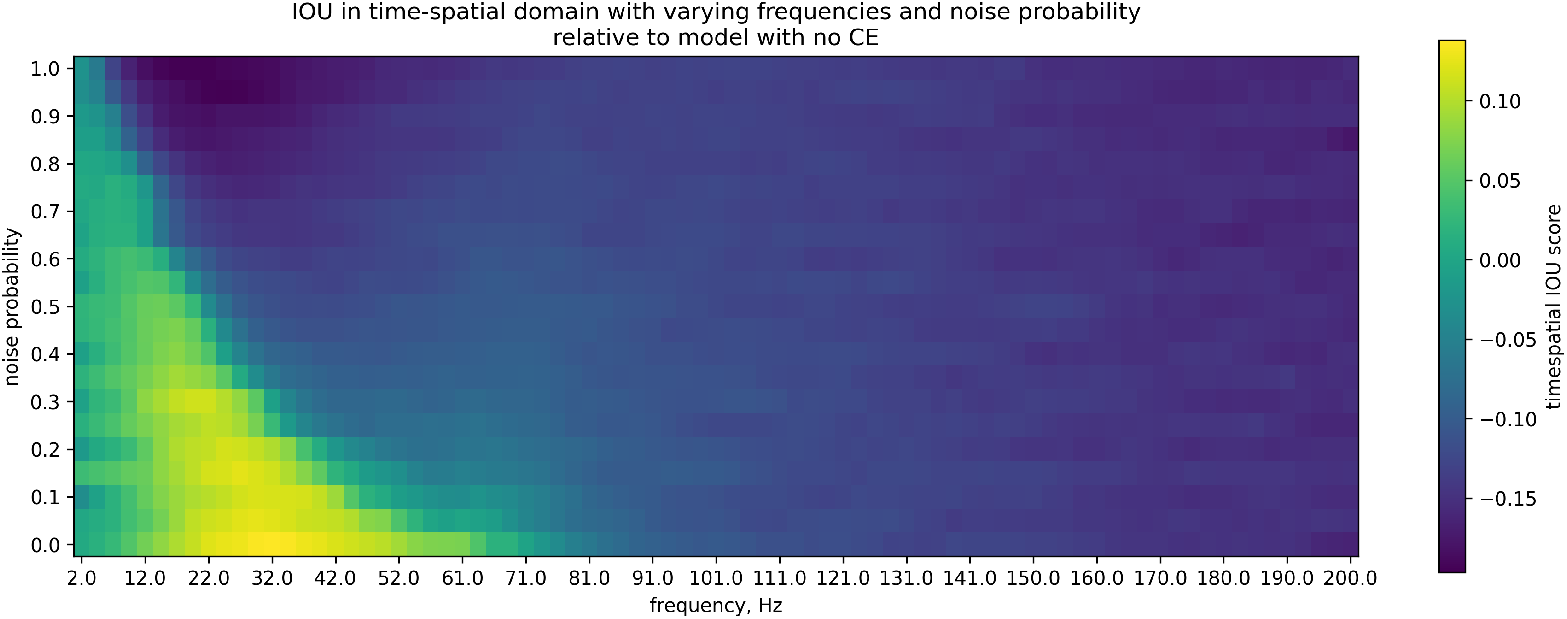
Relative accuracy of the system in time-spatial domain with varying noise probability.

**Fig. 12.**
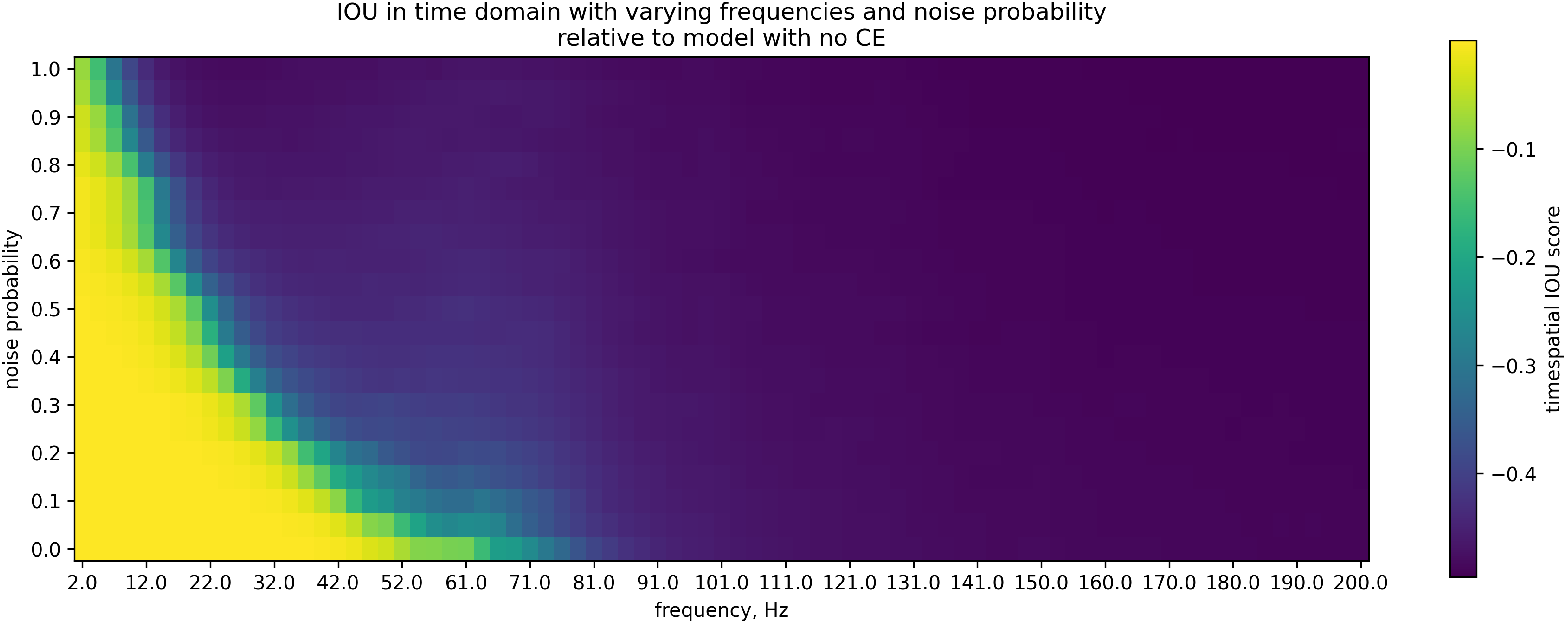
Relative accuracy of the system in time domain with varying noise probability.

**Fig. 13.**
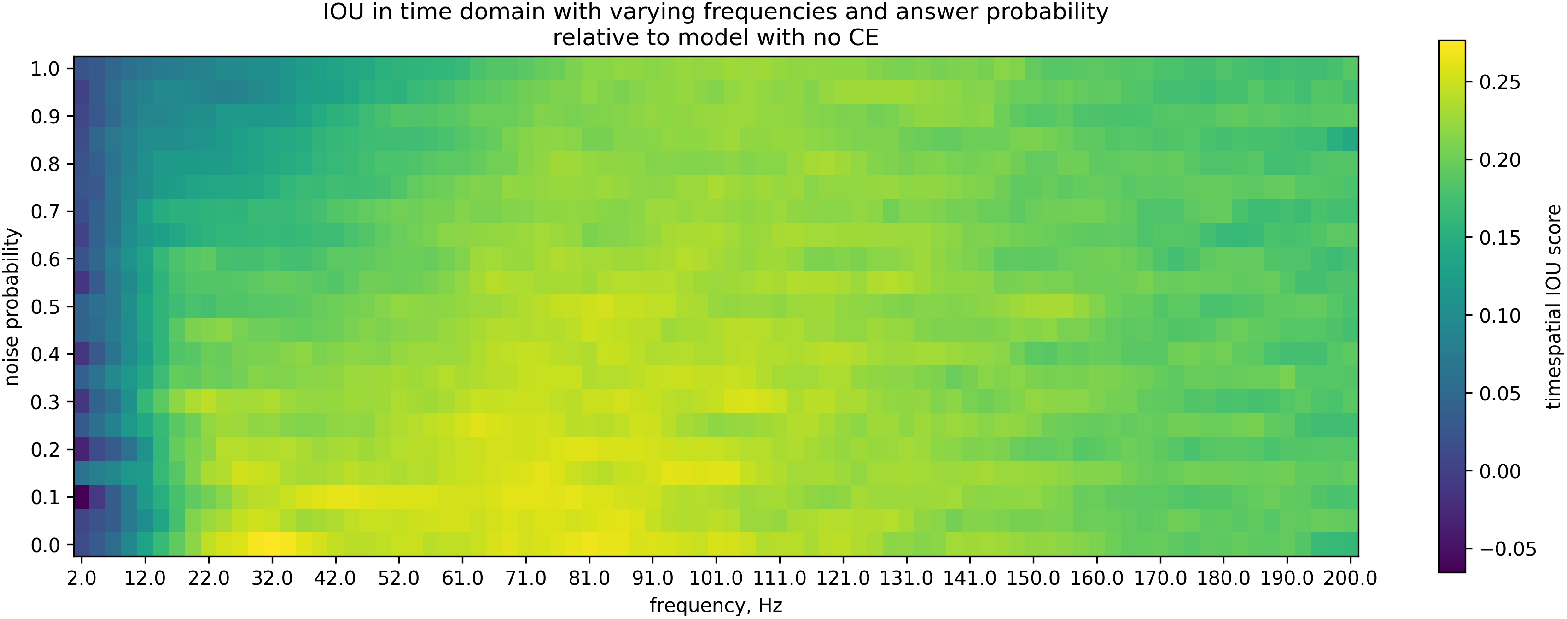
Relative accuracy of the system in spatial domain with varying noise probability.

**Fig. 14.**
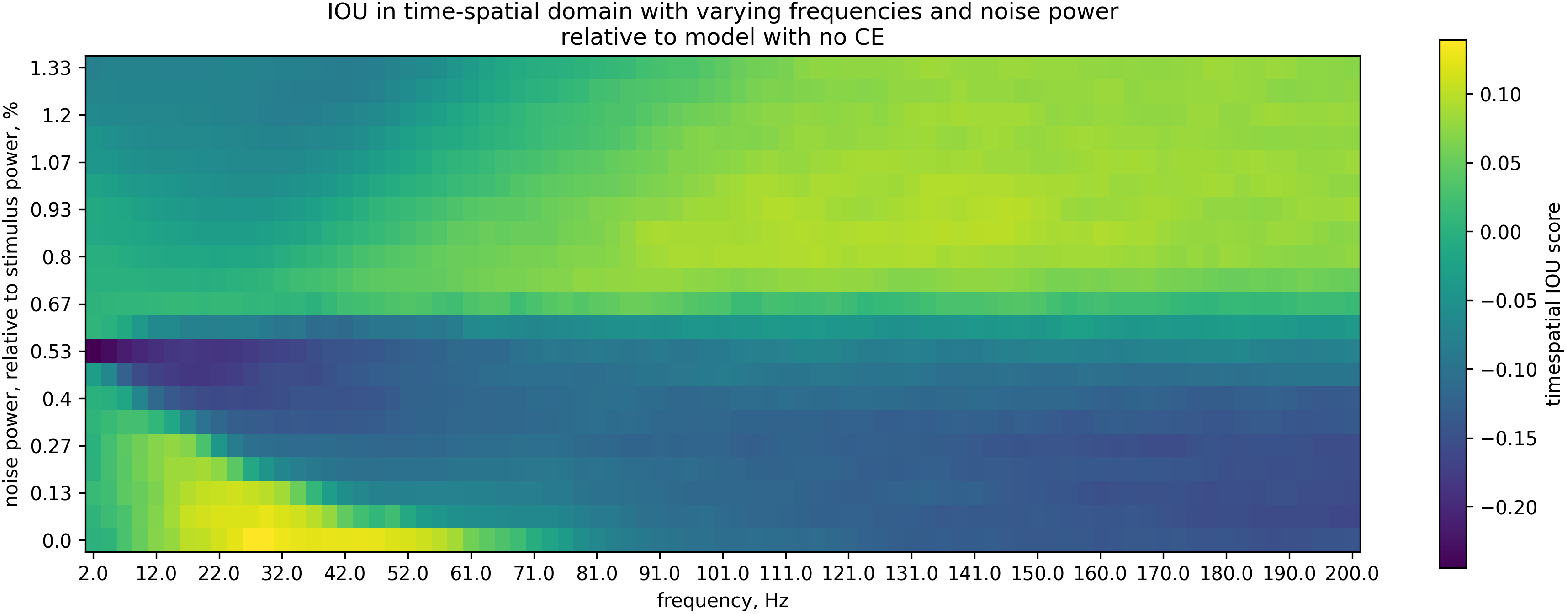
Relative accuracy of the system in time-spatial domain with varying noise amplitude.

**Fig. 15.**
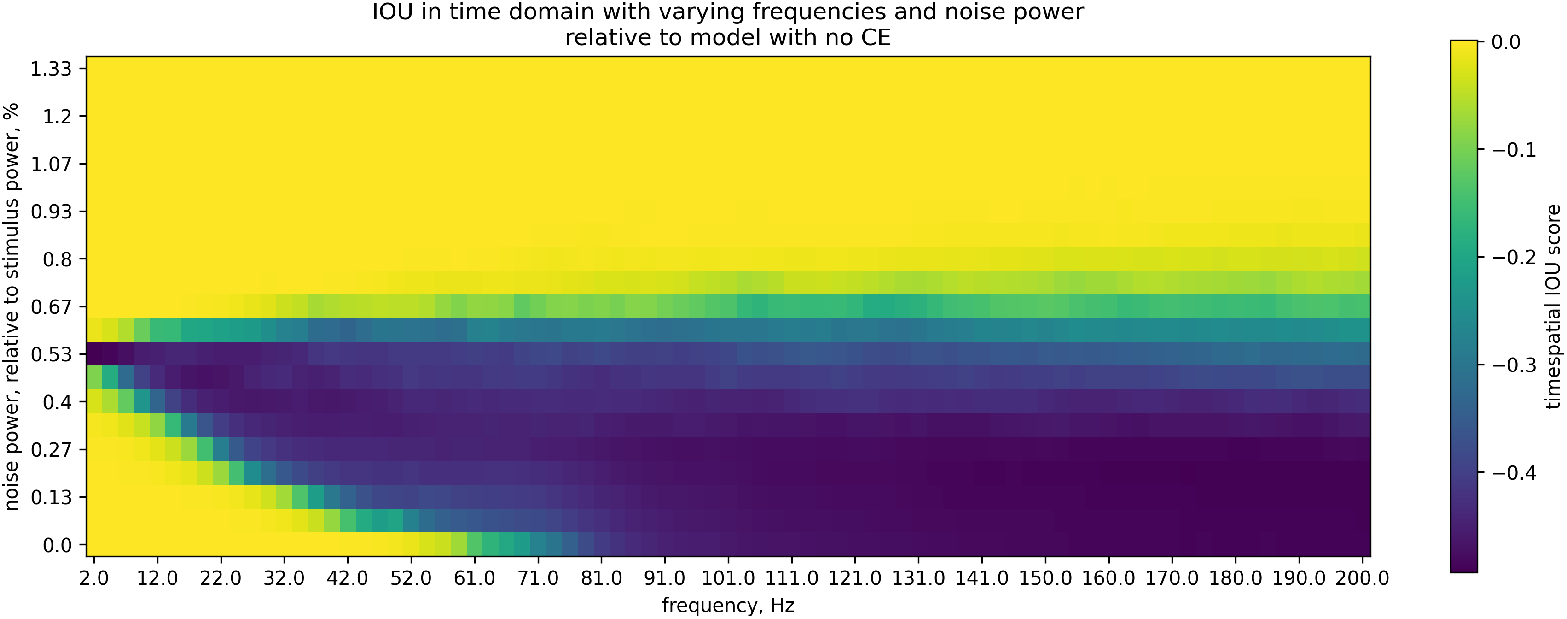
Relative accuracy of the system in time domain with varying noise amplitude.

**Fig. 16.**
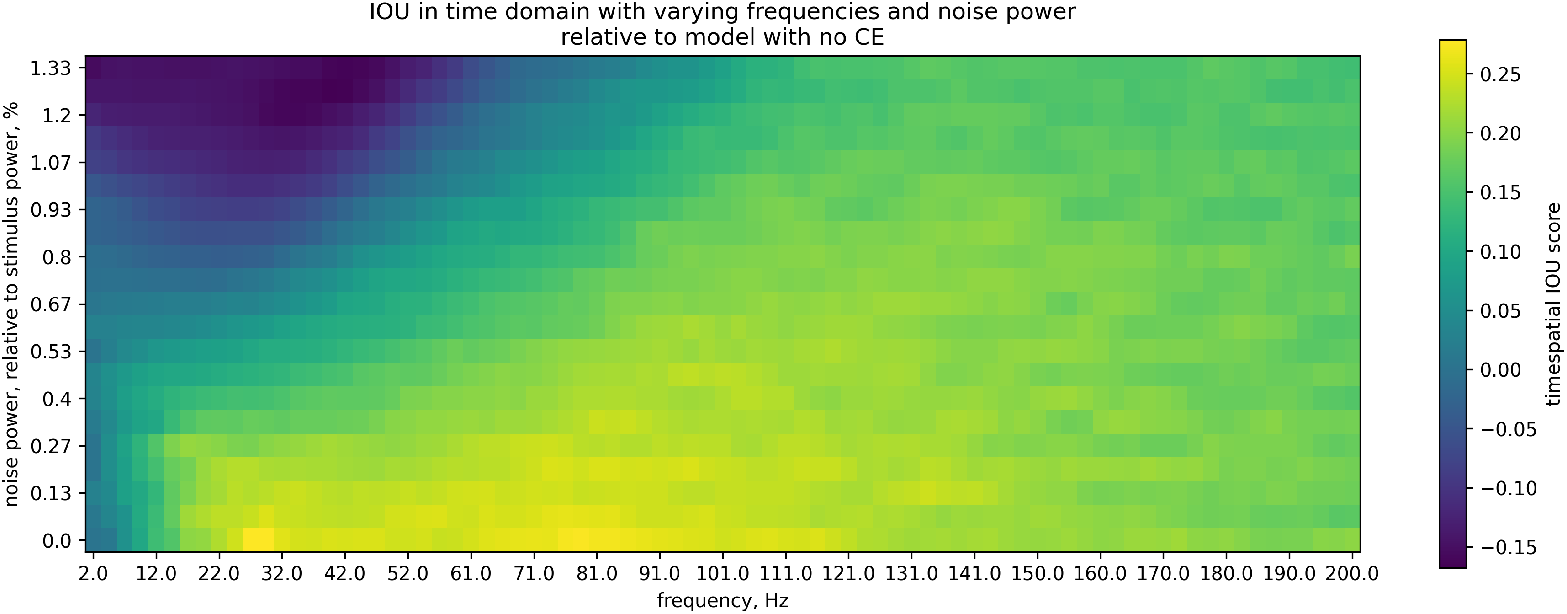
Relative accuracy of the system in spatial domain with varying noise amplitude.

### 3.3 Experiment 3

To evaluate network’s sensitivity to the density of afferent stimulation, the stimulation probability parameter was systematically varied. From a physiological perspective, this parameter represents the fiber recruitment ratio (Dalrymple et al. 2021) — a fraction of peripheral axons within a nerve that has successfully transmitted an action potential to the DRG somata during a particular stimulation event. The relative gains of the network are shown in Fig 18; the median gain in the temporal domain was negative (*≈ −*0.0916 *±* 0.023; Fig 19), as opposed to the gain in the spatial domain (median *≈* 0.0395 *±* 0.019; Fig 20).

**Fig. 17.**
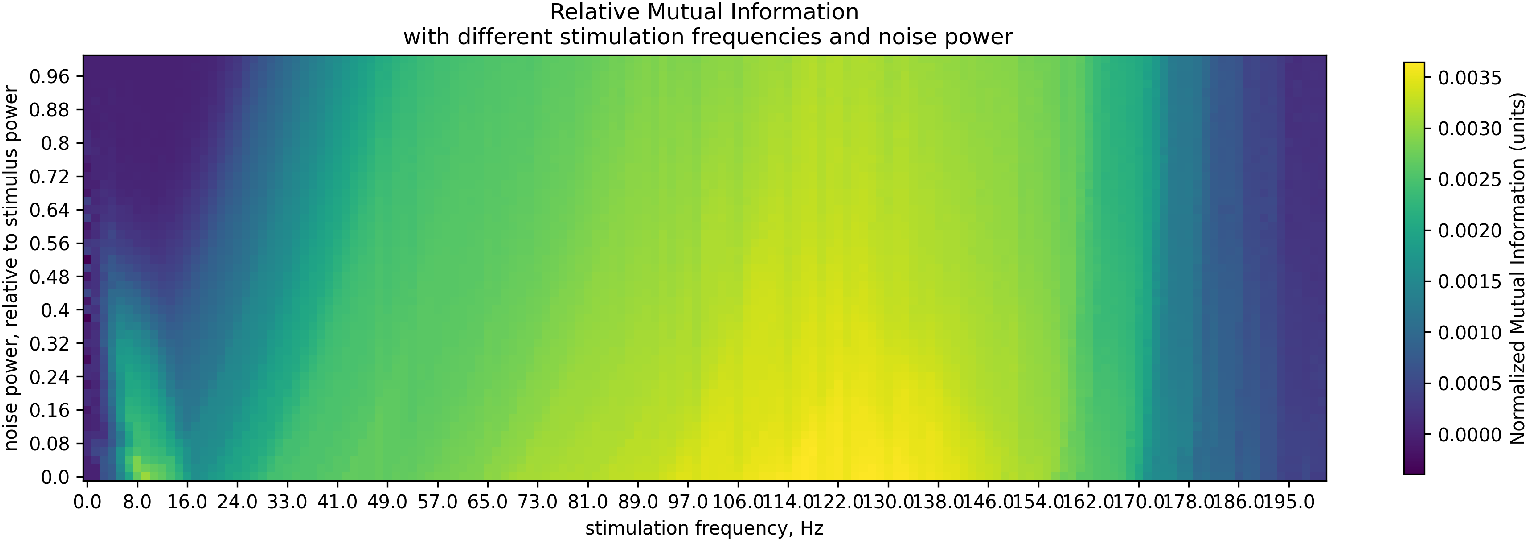
Relative mutual information under different stimulation conditions and varying noise power.

**Fig. 18.**
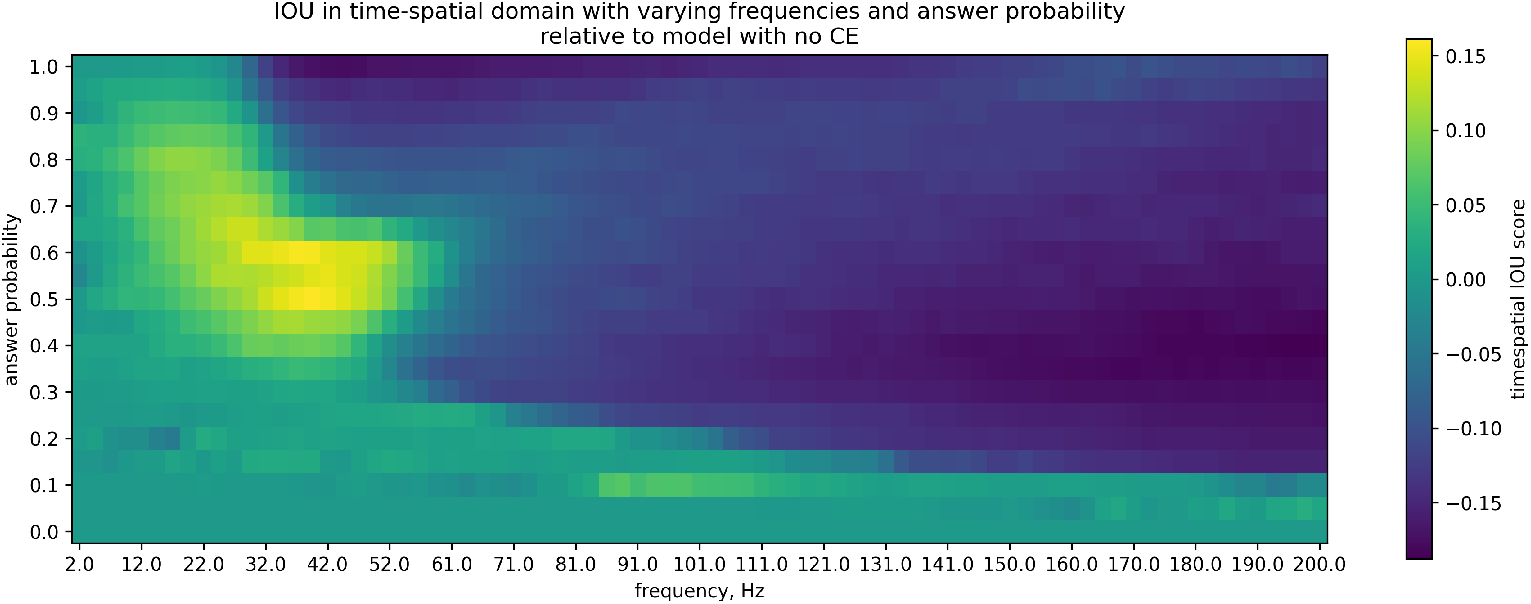
Relative accuracy of the system in time-spatial domain with varying afferent stimulation density.

**Fig. 19.**
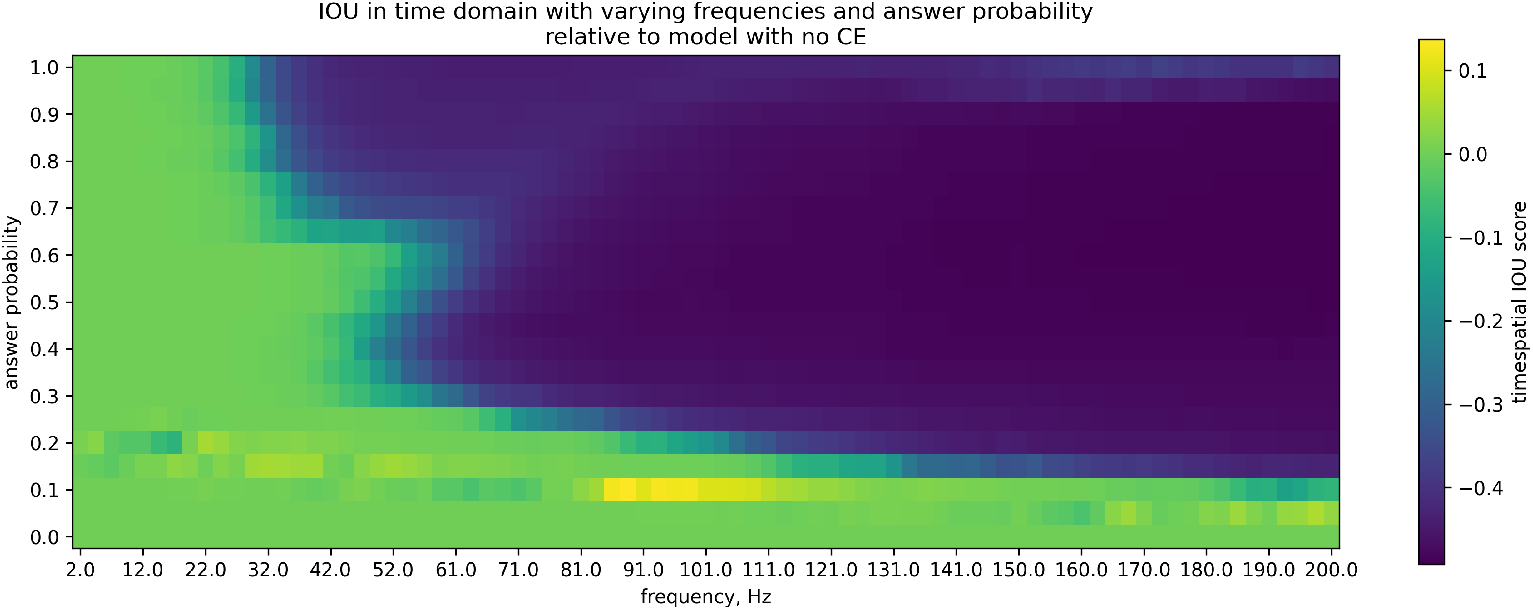
Relative accuracy of the system in time domain with varying afferent stimulation density.

**Fig. 20.**
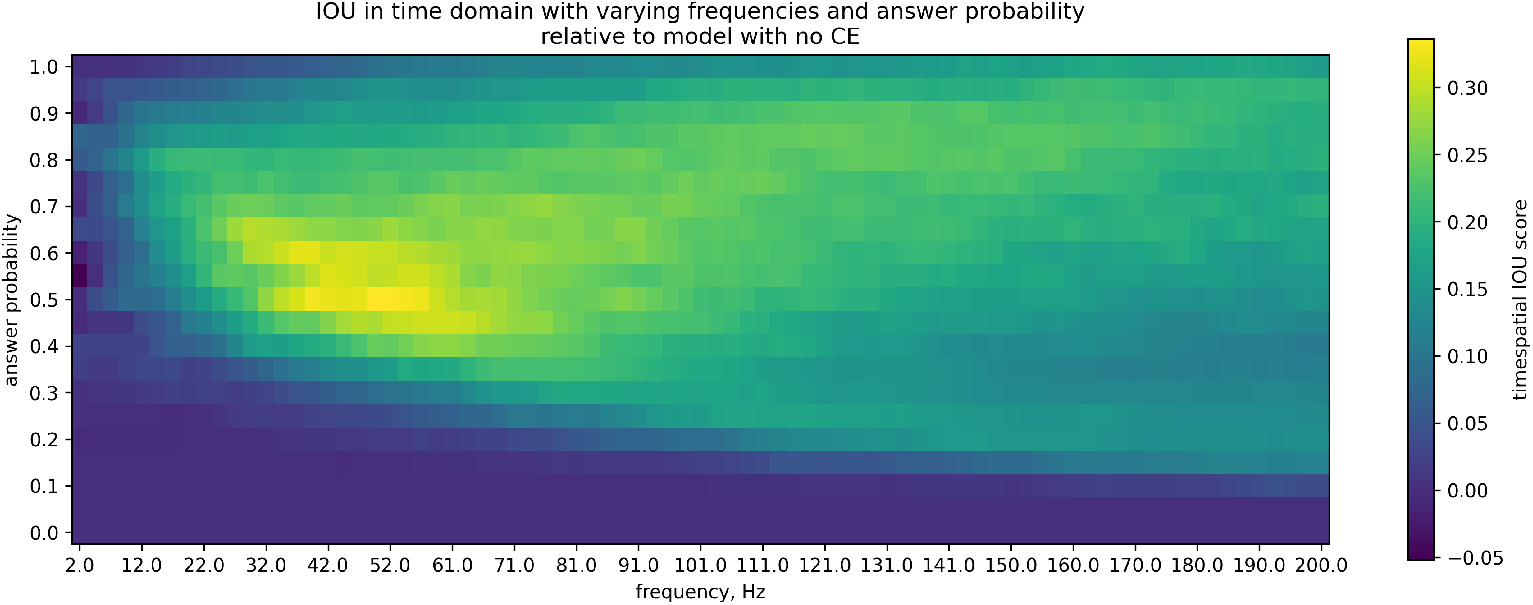
Relative accuracy of the system in spatial domain with varying afferent stimulation density.

### 3.4 Experiment 4

The most significant assumption of our model is that neighboring neurons always innervate neighboring receptive fields (i.e., strict somatotopy). This is a significant simplification: whilst it could be true for most of the neurons, some (still unknown) proportion can be shuffled. Hence, we introduced a degree of distortion metric — after a model is generated, the cross-excitation matrix is mixed, effectively swapping a portion of neurons between each other. This way we obtain a network in which a chosen proportion of neurons are not following somatotopy. The aim is to test whether the positive effects of DRG (physiological utility) require strict somatotopy. Fig 21, Fig 22 and Fig 23 demonstrate the performance of the network under varying degree of distortion (DoD). We observe familiar dynamics — while the system is under a very low stimulation frequency, the gains of cross-excitation are the most prominent. The score rapidly decreases with stimulation frequency, and, at high frequencies, both the DoD and CE presence becomes more disruptive. However, the clear spatial gains are present in most DoD-frequency combinations — the median spatial gain was *≈*0.03685 *±*0.0095 and the temporal gain was *≈−*0.435*±*0.137 (a very significant degradation) with the spatio-temporal gain of *≈−*0.2*±*0.05. Overall, the spatial IOU had a clear negative trend with DoD (Fig 24), which is expected for spatial accuracy, while the temporal score did not change with DoD (Fig 25).

**Fig. 21.**
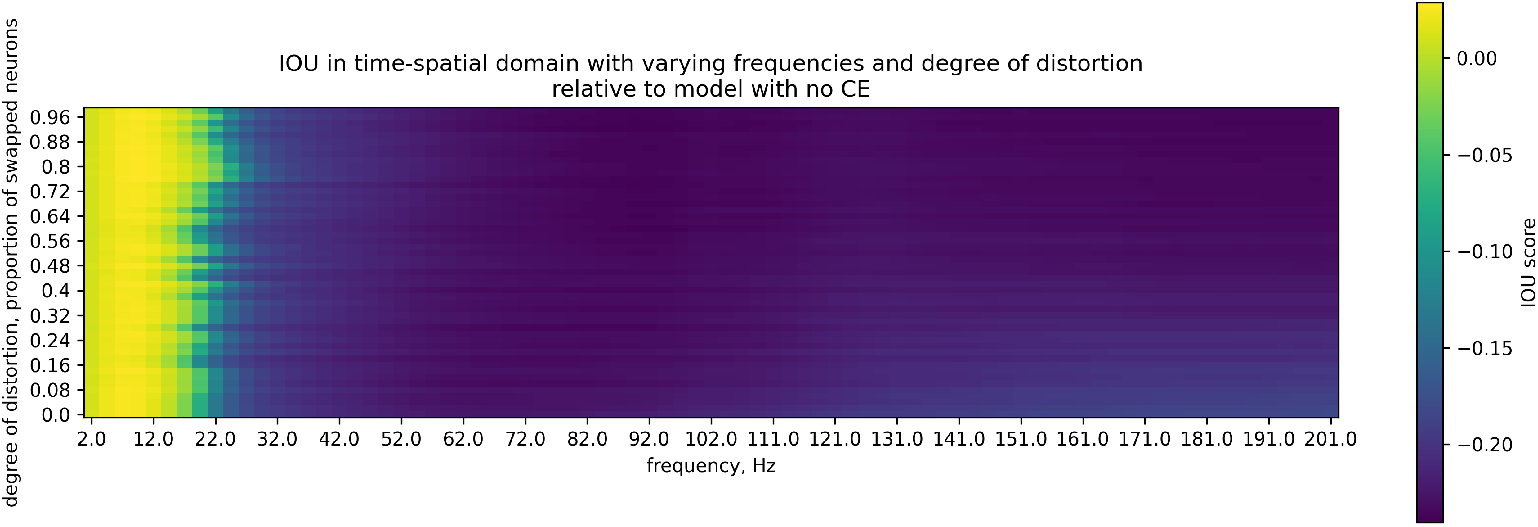
Relative accuracy of the system in time-spatial domain with varying frequency and degree of distortion.

**Fig. 22.**
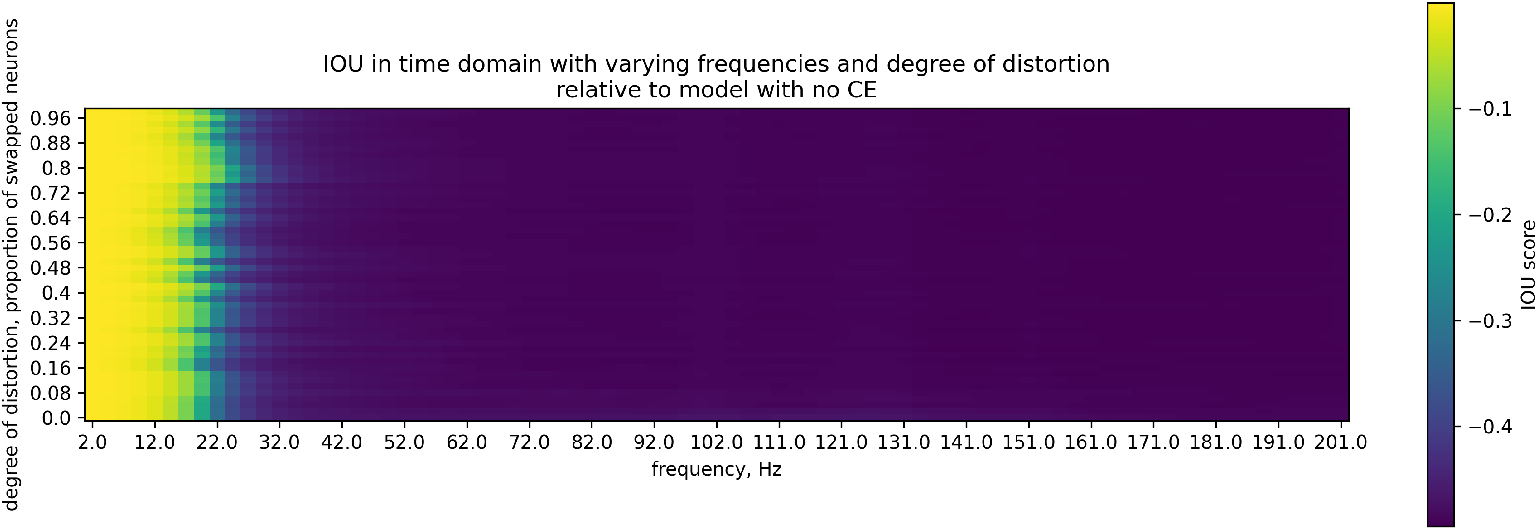
Relative accuracy of the system in time domain with varying frequency and degree of distortion.

**Fig. 23.**
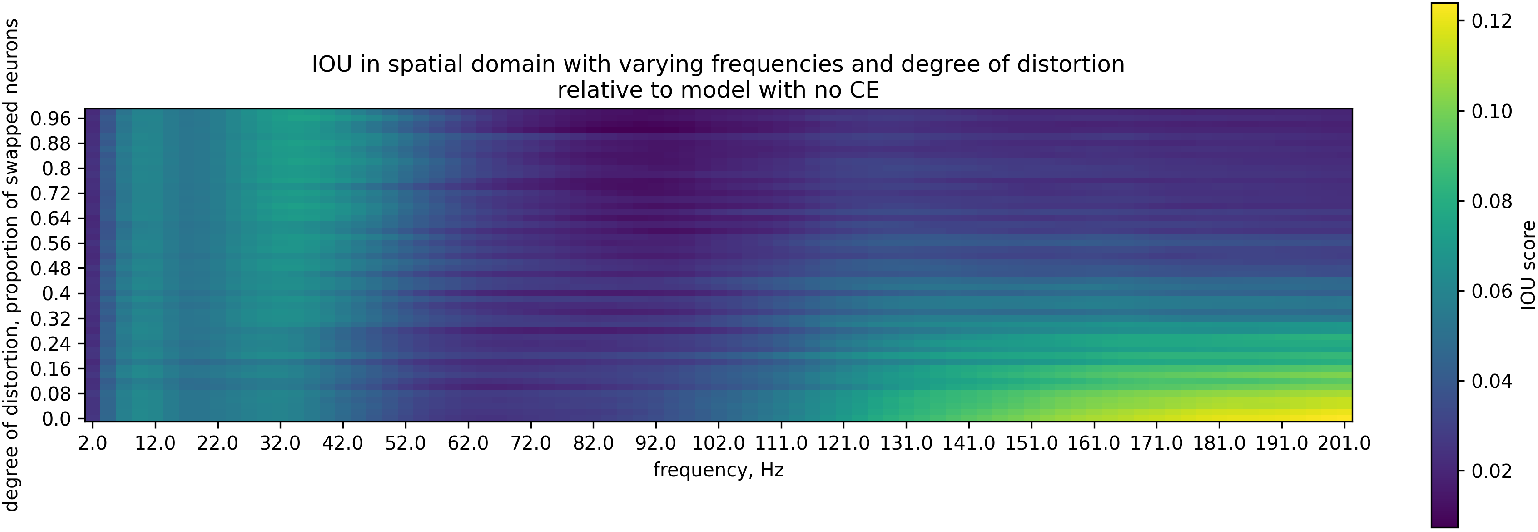
Relative accuracy of the system in spatial domain with varying frequency and degree of distortion.

**Fig. 24.**
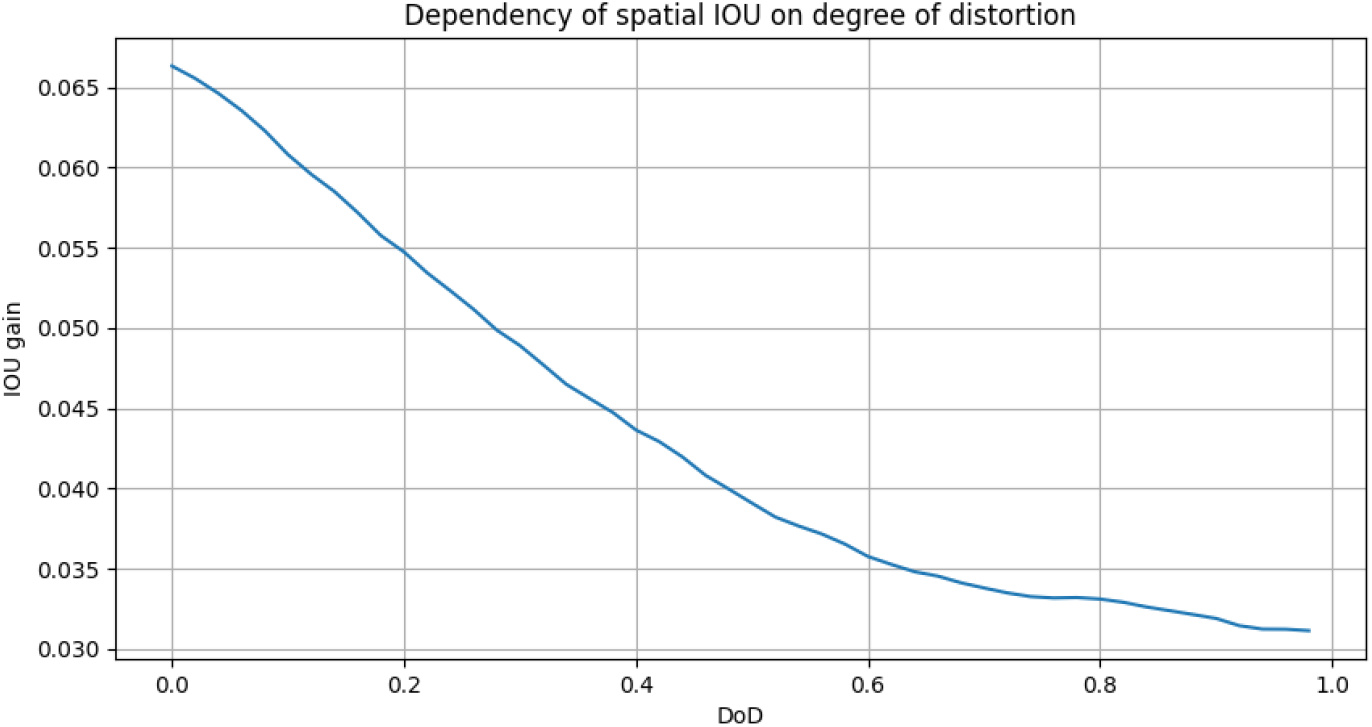
Dependence of spatial IOU score on degree of distortion.

**Fig. 25.**
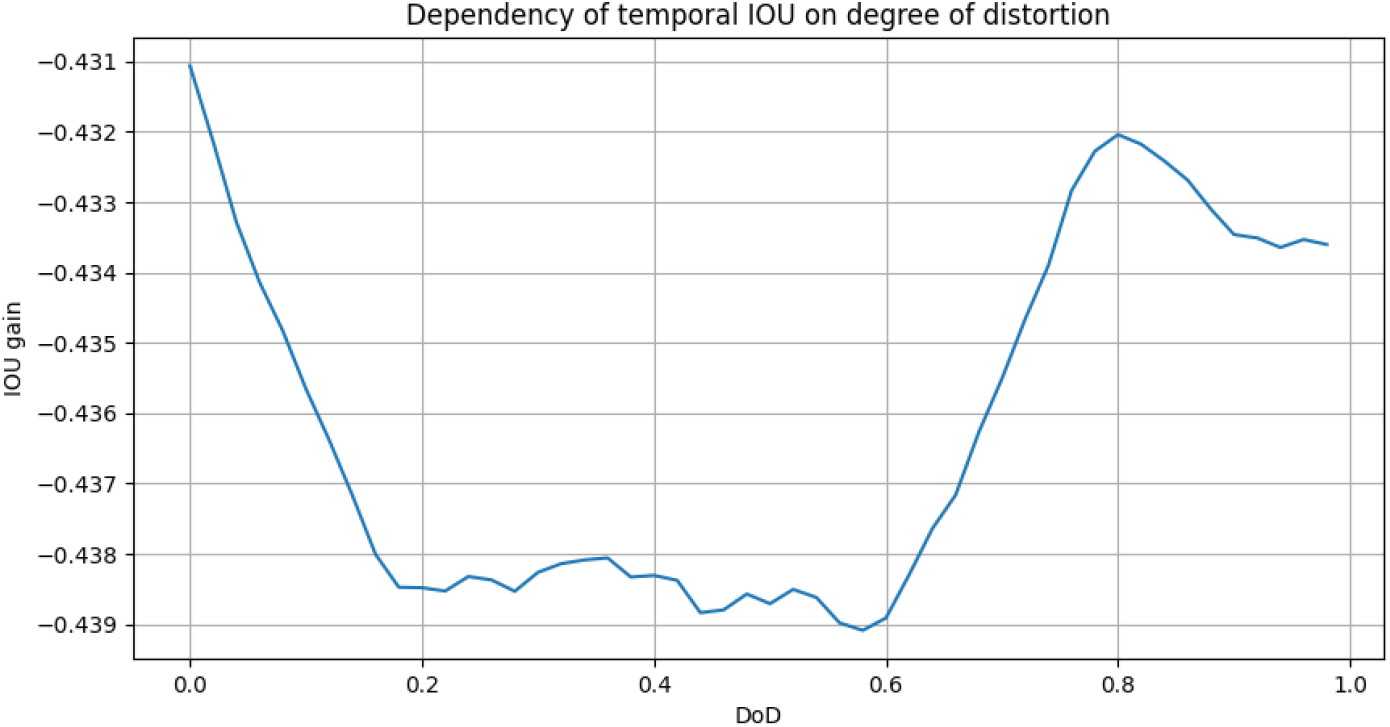
Dependence of temporal IOU score on degree of distortion.

## 4 Overall results

The derivations above make four falsifiable claims about a network of *N* = 400 LIF neurons coupled only by a diffusible agent (*τ* = 220 ms, *g*_*ce*_ = 0.22). By the chain rule (10), the population response splits into a spatial term *I*(*L*; *Y* ) and an intensity term *I*(*A*; *Y*| *L*), which our spatial-and temporal-domain IOU metrics respectively track. The claims are: (i) cross-excitation (CE) trades the second for the first; (ii) the cost is a fixed number of bits set by the coupling constant *K* and independent of the stimulus (13); (iii) the benefit is regime-dependent through the product (1*− p*_0_) *p*_CE_ *ρ*_soma_ (14); and (iv) the net exchange (15) is close to break-even in raw bits. We test each in turn. All gains are differences against a CE-free network run under identical stimulation.

### 4.1 The composite metric masks the exchange

Absolute accuracy was dominated by stimulation frequency, not by the coupling, under both ideal (Fig 2) and noisy (Fig 3) conditions. Referenced against the uncoupled network, the composite spatio-temporal score showed no meaningful benefit: the maximum gain across the entire frequency–range space was 0.022 (ideal) and 0.00115 (noisy), with optimal ranges of only 3 and 1 affected neighbours (Fig 4, Fig 5).

This null is a cancellation. Decomposed, the temporal component never gained significantly (maxima *≈ −*2.59*×*10^*−*6^ ideal, 0.0002 noisy; Fig 6, Fig 7) while the spatial component gained substantially ( *≈*0.12 ideal, *≈*0.2264 noisy; Fig 8, Fig 9), optimizing at much wider ranges of 7 and 13 neighbors. Prediction (i) holds: the two terms of (15) move in opposite directions, and the composite metric is the sum of a cost and a benefit rather than a measure of either.

### 4.2 The temporal cost is fixed, as (13) requires

Because the CE-induced variance 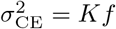 (11) grows in *f* in the same proportion as the mean drive 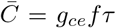 (7), the frequency cancels from the effective SNR and the intensity loss collapses to 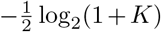 — a constant. The prediction is specific: the cost should be flat along the frequency axis, should scale with *N*_*k*_ (which enters *K* linearly), and should be blind to anything not appearing in *K*.

All three signatures are present. The relative temporal maps (Fig 6, Fig 7) show degradation that is essentially uniform across 2–200 Hz and organised instead by interaction range, with near-zero cost confined to a narrow band of small ranges before saturating at *≈−*0.4 beyond it. The sharpest test is somatotopic distortion: *ρ*_soma_ does not appear in (13), so shuffling the interaction matrix should leave the cost untouched. It does — the temporal gain varies only between *−*0.431 and *−*0.439 across the full range of distortion (Fig 25), a variation of *<* 2% against a spatial term that halves over the same sweep. The cost of CE is a property of the coupling, not of what the coupling is wired to.

### 4.3 The spatial benefit

Equation (14) predicts a benefit that is single-peaked in frequency: *p*_CE_ switches on near *f*_th_ (9) and saturates, while the room for improvement (1 *− p*_0_) shrinks as stimulation makes the baseline code reliable. Each factor is separately manipulable, and each sweep isolates one.

#### Frequency

Spatial gains concentrate at low-to-moderate frequency and fade toward the high-frequency edge (Fig 9, Fig 8), where the uncoupled code is near-saturated and few gaps remain to fill. Since *f*_th_*∝* 1*/N*_*k*_, the wide optimal ranges found above (7– 13 neighbours) are what a mechanism gated by accumulation should select: broader coupling lowers the switch-on frequency into the operating band.

#### Noise

Under noise the baseline is unreliable, (1 *p*_0_) is large, and the maximal spatial gain nearly doubles ( *≈*0.2264 noisy vs *≈*0.12 ideal). Sweeping noise occurrence and amplitude at fixed range 13 reproduces the exchange with stable polarity — median spatial gains of *≈*+0.054 (Fig 13) and *≈*+0.0454 (Fig 16) against median temporal costs of *≈−* 0.12 (Fig 12) and *≈−*0.0555 (Fig 15); composites in Fig 11 and Fig 14. CE helps most exactly where the system is stressed.

#### Recruitment

Varying the fiber recruitment ratio drives both factors at once and therefore provides the strongest test of (14): the benefit should vanish at both extremes. It does. Median gains preserve the exchange ( *≈*+0.0395 spatial, Fig 20; *≈−*0.0916 temporal, Fig 19), but the gain is strongly non-monotonic, peaking at intermediate recruitment (*≈*0.4–0.7) and moderate frequency (Fig 18) — complete afferent images leave nothing to recover, and sparse ones generate too little agent to recruit neighbours.

#### Somatotopy

Shuffling the interaction matrix should erode the benefit monotonically while, as shown above, leaving the cost fixed. The spatial gain falls steadily from *≈*0.066 undistorted to *≈*0.031 at near-complete shuffling (Fig 24), with the exchange persisting but the composite driven to *≈−*0.2 across the sweep (Fig 21–Fig 24). Roughly half the benefit survives full randomisation, indicating that part of the gain is generic sub-threshold recruitment while the somatotopy-gated part is the half that (14) describes.

### 4.4 The net exchange is near-neutral in raw bits

Equation (15) predicts a positive balance where gap-filling dominates — low-to-moderate frequency, modest range, noisy conditions — turning mildly negative where the fixed cost dominates. The relative mutual-information map (Fig 10) has exactly this geometry, positive at low-to-moderate frequency and small range and shading slightly negative at high frequency and wide range. The pattern recurs under noise amplitude (Fig 17) and distortion (Fig 26), the latter turning negative precisely where *ρ*_soma_ is degraded and frequency is high. CE is thus close to information-neutral on a bit-for-bit accounting, with a small positive balance in most conditions — it redistributes capacity rather than creating it.

**Fig. 26.**
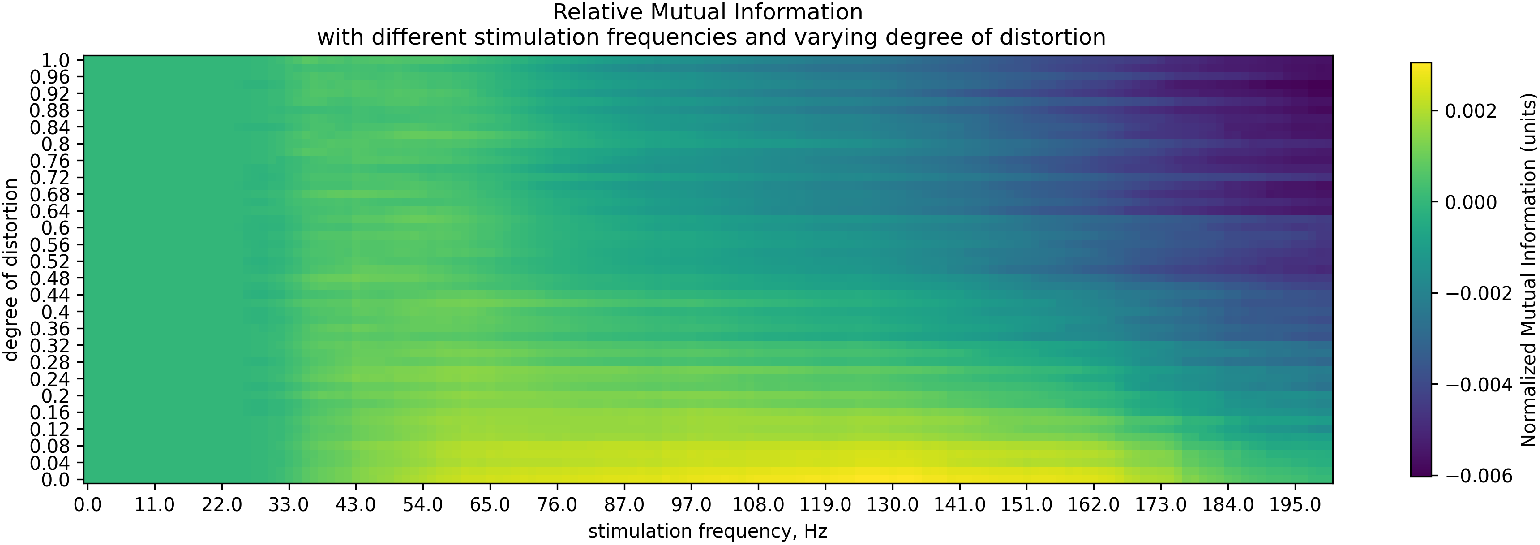
Relative mutual information under different stimulation conditions and varying degree of distortion.

### 4.5 Weighted by perceptual value, the exchange is favorable

Raw bits weight a spatial and an intensity bit equally. The utilization fraction (18), *U* (*f* ) *∝*log_2_(ln *f* )*/f*, falls monotonically for *f > e*: the rate channel’s capacity grows linearly with *f* while the number of perceptually distinguishable intensity levels grows only logarithmically, so that only *≈* 10% as much of the channel is perceptually usable at 200 Hz as at 10 Hz. The bits CE spends are drawn from this surplus. The consequence is visible as an asymmetry across the operating range rather than as a single number: where the direct spatial payoff is largest (low-to-moderate frequency, noise, partial recruitment) CE helps materially, and where the payoff is smallest (high frequency, saturated code) the intensity it corrupts is precisely the intensity that perception cannot resolve. The exchange that reads as break-even in Fig 10 is therefore favourable once weighted: CE spends capacity that is cheap and getting cheaper to buy capacity that stays valuable.

## 5 Model limitations

As with any in silico model, the presented hybrid approach relies on several simplifying assumptions. The most significant of these is the assumed kinetics of cross-excitation (CE). We implemented a simplified, mediator-based mechanism as the sole driver of CE, though it could be argued that this signaling might also occur via simple ionic diffusion or specific neuron-glia interactions (or interplay of those). Furthermore, our model assumes that the signaling agent is transmitted instantaneously and depolarizes neurons linearly as a function of its concentration. Both of these assumptions are defensible provided that CE connects strictly to immediately adjacent neurons – consistent with how the phenomenon was originally discovered experimentally. Additionally, an in vivo mediator would likely show more complex dynamics, such as context-dependent effects or the presence of compensatory homeostatic mechanisms, which we also neglect here. However, the primary aim of the present work was not to construct an exact biophysical model of the – as yet incompletely understood – interactions between dorsal root ganglion (DRG) neurons. We adopted these simplifications because our objective was to analyze the overall direction and functional utility of the CE effect, rather than to determine its precise biophysical parameters. By abstracting the micro-level details, our model successfully isolates the systemic consequences of CE on the network. Ultimately, we wanted to outline a macroscopic perspective of *what* cross-excitation could do to a neural signal, and *what utilities* it brings to a system as a whole.

## 6 Discussion

The central result — that an excitatory, spatially diffuse coupling improves spatial accuracy — is counter-intuitive, because the canonical circuit motif for improving localization is lateral inhibition, which sharpens the population image by suppressing off-peak activity. CE does the opposite operation and nevertheless improves the spatial IOU. The two are not in conflict because they decrease different types of errors. Lateral inhibition removes false positives at the edges of an already complete activity profile; CE removes false negatives inside an incomplete one. In our model the uncoupled population image is punctate and incomplete: sub-threshold afferent drive, stochastic recruitment failure, and heterogeneous thresholds leave silent neurons inside the stimulated territory. CE fills exactly those gaps, because the agent concentration at a given soma is proportional to the recent firing of its somatotopic neighbors and itself, i.e., it is highest precisely where the stimulus already is. This also explains why the benefit is gated by somatotopy (Fig 24) and why roughly half of it survives full shuffling: the shuffled component is generic sub-threshold recruitment (any silent neuron is easier to fire), while the somatotopy-dependent component is the spatially structured completion that the intact wiring provides.

Gap-filling is the most visible consequence of cross-excitation, but it is not the operation itself. What the model implements is more elementary: the agent concentration at soma is a weighted sum of the recent spiking of its somatotopic neighbors, Eq (7), so the population converts the sharp afferent image into a smoothed field, and only then applies the spiking non-linearity. CE is therefore a convolution — spatial kernel of half-width *N*_*k*_, temporal kernel of width *τ* — inserted before the thresholding stage rather than after it. Gap-filling is what this convolution looks like when the image is sparse but it is not the only thing the convolution does. Three consequences follow that gap-filling alone does not cover. Because the input summed at a soma is smeared over *N*_*k*_ neighbors, a stimulus too weak to bring any single neuron to threshold can still be detected if it covers enough of them. The population becomes a filter for spatial extent: coupling therefore trades single-unit sensitivity for a lowered detection threshold on contiguous stimuli. This is distinct from filling gaps in an already supra-threshold response, and it predicts that CE should lower absolute detection thresholds for broad, low-amplitude contacts more than for punctate ones. We did not test this directly. Because the kernel is positive and (at least somewhat) somatotopically arranged, CE preferentially completes activity patterns that are already spatially contiguous, and does nothing for isolated spikes, whose deposited agent has no coincident neighbors to sum with. The coupling thus encodes, in hardware and without any learned parameters, the statistics of natural tactile input: real contacts are usually connected patches of skin (sensed by several neighboring neurons), whereas noise is spatially independent. Isolated noise spikes are not amplified into false population responses because a single neighbor’s contribution is small relative to threshold but coherent activity is. Somatotopic distortion degrades the benefit because it breaks the match between a stimulus statistic and the kernel’s assumptions. The same argument applied along the time axis explains the temporal penalty. With *τ* = 220 ms the agent is a leaky integrator of recent population history, so the smoothing is spatiotemporal, not purely spatial. Smoothing in time is what corrupts onset and offset boundaries and inflates the false-positive count during gap periods. The cost and the benefit are not two mechanisms but one operation projected onto two axes: the kernel that completes the spatial image also blurs the temporal one. Finally, the somatotopy-independent half of the gain (Fig 24, *≈*0.031 surviving near-complete shuffling) is naturally read in this framework as the DC component of the filter — a uniform, activity-dependent excitability increase that raises the population’s overall responsiveness in proportion to how much of it is currently firing.

### 6.1 An analogy from machine vision

The pairing of a spatially diffuse excitation with a downstream inhibitory surround has a direct counterpart in image processing, where derivative operators are never applied to a raw noisy image: edge-detection pipelines low-pass filter first and differentiate second, and the two steps are routinely composed into a single centre-surround kernel. The somatosensory pathway may separate these stages anatomically. The ganglion supplies the excitatory, isotropic smoothing; the dorsal column nuclei supply the lateral inhibition. Therefore, CE can be interpreted as an analogy for a classical CV preprocessing step.

## 7 Conclusion

We asked whether cross-excitation in the dorsal root ganglion is an imperfection of somatic isolation or a functional component of somatosensory pre-processing. Simulating a population of 400 LIF neurons coupled solely by a diffusible agent calibrated to reported CE parameters, and analyzing the network with a decomposed spatio-temporal IOU metric together with mutual information, we find that CE is neither noise nor a general-purpose improvement. It is a redistribution.

The composite accuracy metric shows essentially no effect of CE, but this null is a cancellation of two opposing terms. Temporal (intensity) accuracy is degraded by an amount that is flat across 2–200 Hz, scales with interaction range, and is insensitive to somatotopic shuffling — the signature of a fixed cost set by the coupling constant alone. Spatial accuracy, by contrast, is improved by an amount that is strongly regime-dependent: single-peaked in frequency, roughly doubled under noise, maximal at intermediate fiber recruitment, and monotonically eroded by loss of somatotopy. The two behaviors follow directly from the model equations, which predict a mean CE drive linear in firing rate and neighbor number, a switch-on frequency proportional to 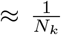, and a benefit depending on spiking frequency and strictness of somatotopy. On a bit-for-bit accounting the exchange is close to neutral, with a small positive balance at low-to-moderate frequency, modest coupling range, and degraded input quality. That accounting, however, weights an intensity bit and a spatial bit equally. Because rate coding supplies capacity linearly in firing rate while perceived intensity grows only logarithmically (Weber–Fechner), the fraction of the rate channel that intensity discrimination can perceptually use falls monotonically with stimulus strength. The bits CE spends are drawn from this growing surplus, whereas the bits it buys — spatial location — are perceived without compression. The trade is there-fore perceptually favorable precisely where it looks least profitable in raw bits. Three consequences follow. First, the DRG is better described as an active, chemically mediated spatial pre-filter than as a passive relay with imperfect insulation. Second, CE is most useful when the afferent code is unreliable — under noise or partial recruitment —which offers a functional account of its near-absence in nociceptive C-type neurons, where smearing across neurons carries a cost with no corresponding localisation benefit. Third, the mechanism is gated by somatotopy: it injects structured rather than random perturbation, and roughly half its benefit disappears when neighbour relations are randomised. These conclusions rest on an abstracted mediator model and remain to be tested experimentally. The framework nevertheless yields concrete, falsifiable predictions — a stimulus-independent intensity cost, a coupling-range-dependent switch-on frequency, and a two-point discrimination advantage that should be largest under noisy or weakly recruited afferent input — each accessible to psychophysical or electrophysiological tests.

## Statements and Declarations

### Data&Code availability

All the code will be made available at our GitHub repository.

### Author contributions

D.P. - conceptualization, methodology, software, writing - orginal draft; N.K. - conceptualization, formal analysis, writing - review & editing.P.M. - writing - review & editing, supervision, funding.

